# Spreading alpha-synuclein Oligomers Trigger Astrocyte Changes and Astrocyte-Glutamatergic Neuron system dysfunction in an Age-related Manner

**DOI:** 10.1101/2025.08.08.669249

**Authors:** Jaehyun LeeBae, Verena Bopp, Alicia Goreth, Veselin Grozdanov, Julia K. Kühlwein, Ermanno Gazzola, Daniel Rombach, Laura Meier, Leda Dimou, Karin M. Danzer

## Abstract

**Background:** Parkinson’s disease (PD) is characterized by the progressive accumulation and spatio-temporal spread of α-synuclein (α-syn) oligomers and a progressive loss of dopaminergic neurons. Many studies showed a direct cytotoxic effect of α-syn oligomers on neurons. Other cell types including astrocytes were also reported to show specific responses to α-syn and are believed to play a role in the spreading of PD pathology.

**Methods:** To investigate the transcriptional and cellular consequences of α-syn oligomer spreading, we employed spatial transcriptomics and single-nucleus RNA sequencing (snRNA-seq) in a transgenic PD mouse model expressing human α-syn in neurons. We further compared our findings to published public snRNA-seq datasets from human PD patients

**Results:** Our analysis identified α-syn spreading mostly to the substantia nigra and defined a transcriptional “Spreading Signature” associated with α-syn pathology. We found an age correlated increase in astrocytes, close interactions between astrocytes and α-syn, and transcriptional dysregulation of the astrocyte-glutamatergic neuron axis. We further identified two subtypes of glutamatergic neurons that are vulnerable to astrocytic changes. Comparative analysis with human PD snRNA-seq data showed concordant transcriptional changes related to astrocytic dysfunctions and diminished neuronal signaling.

**Conclusion:** Based on our results, we propose a model of α-syn oligomer spreading involving astrocytes, glutamatergic synapses, and a disturbance in the astrocyte-glutamatergic neuron axis.

## Background

Parkinson’s disease (PD), a chronic progressive neurodegenerative disease, is characterized pathologically by loss of dopaminergic neurons in the substantia nigra (SN) and the accumulation of aggregated α-synuclein (α-syn) in the form of Lewy bodies(1). Histopathological postmortem studies suggest that the α-syn pathology starts in the olfactory bulb and brainstem and then spreads progressively to axonally connected areas throughout the CNS in a spatio-temporal manner, eventually reaching the SN(2). Subsequent studies supported this theory as they reported the propagation of α-syn pathology from host to graft in PD patients(3–5) and the cell-to-cell spread of α-syn to anatomically connected regions in PD mouse models(6–9).

A substantial number of studies have demonstrated direct cytotoxic effects of pathologic α-syn on neurons, e.g. by pore formation and calcium influx, sequestering of the Na^+^/K^+^-ATPase, oxidative stress and mitochondrial damage(10–12). However, other cell types can also be affected: microglia, astrocytes, oligodendrocytes and infiltrating immune cells show specific responses to extracellular α-syn and are believed to modulate the spread of pathology(13–15).

A powerful tool to investigate cellular changes by an extrinsic trigger is the study of transcriptomic changes, which can illuminate mechanistic insights and therapeutic targets in affected cells. Recent developments have enabled the resolution of transcriptomic changes to single-cells or spatial domains(16, 17), which might help to elucidate the exact pathological cascade involving α-syn aggregation and transmission.

In a previous study, we established a transgenic PD human α-syn oligomer mouse model, showing neuronal cell-to-cell propagation of α-syn aggregates and α-syn pathology in SN, midbrain, and other regions(18). To investigate the spatial and cell-type-specific effects of α-syn oligomer spreading during aging *in-vivo*, we integrated single-nucleus RNA sequencing (snRNA-seq) with spatial transcriptomics in this model, enabling us to assess the direct consequences of α-syn oligomer propagation.

## Methods

### Generation and Housing of Transgenic Animals

The transgenic mouse model B6.Cg-Tg(Camk2a-tTA)Mmay/DboJ x B6-Tg(ind alpha-syn split Venus)D1 GVO was established in collaboration with Dr. Björn von Einem (Department of Neurology) and Olena Sakk (Core Facility for Transgenic Animals) under the guidance of Dr. Bernd Baumann (Institute of Physiological Chemistry, Ulm University). Detailed phenotypic characterization of these transgenic mice has been previously published(18).

Mice were maintained under standardized environmental conditions at the Animal Research Center of Ulm University. The double-transgenic animals express V1S/SV2 and calcium/calmodulin-dependent protein kinase II alpha (Camk2a)(19), with transgene expression regulated by doxycycline in a Tet-off system(18). To suppress transgene expression, mice received 100 mg/ml doxycycline and 10 g/l glucose in drinking water; to activate expression, water contained glucose alone. Animals were group-housed in open polycarbonate type II long cages under controlled temperature, humidity and a 12-hour light/dark cycle, with food and water *ad libitum*. Cages were enriched with nesting material and polycarbonate shelters.

The V1S/SV2 model employs a bioluminescent protein complementation assay for detecting α-syn aggregation. It expresses human wild-type α-syn fused to either the N-terminal or C-terminal fragment of yellow fluorescent protein (YFP), termed V1S and SV2, respectively. Co-expression of these two fragments enables reconstitution of functional fluorescent protein upon aggregation of α-syn.

### Rotarod experiments

All mice performed accelerating rotarod (Panlab Harvard LE8205) on the same days and in the same order. The experiment was adjusted to increase from 4 to 40 rpm over a period of 300 s and was carried out in three consecutive trials with a 5 min break in between. The latency to fall was recorded and mean values were used for analysis.

### Nuclei Isolation

Approximately 100 mg of brain material was dissected from a single hemisphere (0 to -5 mm relative to Bregma) using a mouse coronal brain matrix (Cell Point Scientific). Tissue from two animals was pooled per preparation. All steps were carried out on ice to maintain sample integrity. Samples were homogenized in 1,400 µl homogenization buffer (320 mM Sucrose, 5 mM CaCl_2_, 3 mM Mg(Ac)_2_, 10 mM Tris HCl pH 8, 0.1 mM EDTA pH 8, 0.1 % Nonidet P40 Substituent (NP-40), 1 mM β-mercaptoethanol, 0.4 U/µl RiboLock in H_2_O) using a douncer (pestles A and B). The homogenate was filtered through 70 µm and 40 µm Flowmi cell strainers to eliminate debris. 700 µl of the filtrate was combined with 450 µl working solution (50 % Opti-Prep, 5 mM CaCl_2_, 3 mM Mg(Ac)_2_, 10 mM Tris HCl pH 8, 0.1 mM EDTA, 1 mM β-mercaptoethanol in H_2_O). A density gradient was prepared using 300 µl of 40 % Opti-Prep, 750 µl of 30 % Opti-Prep and 700 µl of the homogenized tissue mixture. The density gradient was centrifuged at 10,000 g for 5 min at 4 °C. 200 µl of the isolated nuclei was transferred into a 1.5 ml low-DNA-binding tube and mixed with 250 µl 2 % BSA/0.12 U/µl RiboLock in PBS. The mixture was centrifuged twice at 2,000 g for 3 min at 4 °C, resuspending each time in the same buffer. After a final filtration through a 40 µm Flowmi cell strainer, nuclei were pelleted once more under identical conditions. The pellet was resuspended in 50 µl of 1x nuclei buffer (10X Genomics nuclei buffer diluted with 1 mM dithiothreitol (DTT) and 1 U/µl RiboLock in H_2_O). DAPI staining confirmed nuclear membrane integrity before use in snRNA-seq experiments. Procedures were done for three mice per condition.

### Single-nuclei RNA Sequencing

Single-nucleus libraries were prepared using the “Chromium 3’ Single Cell Gene Expression and Library Kit” (10x Genomics) according to the manufacturer’s instructions. 22,000 nuclei from mouse brain tissue were loaded into the Chromium Controller to generate Gel Beads-in-Emulsion (GEMs). Following GEM-RT incubation, cDNA samples underwent recovery, purification and amplification. Quality assessments on the amplified cDNA were conducted using a High Sensitivity DNA Kit (Agilent) on a TapeStation platform.

Libraries were generated through fragmentation, adaptor ligation and Sample Index PCR, then purified and re-evaluated on the TapeStation for fragment quality. Sequencing was performed by Novogene (United Kingdom) on an Illumina NovaSeq 6000.

### Visium spatial brain slice preparation

Fresh-frozen brain tissue was prepared for visium spatial gene expression analysis. Animals were euthanized by fast cervical dislocation and brains were immediately dissected. A metal beaker of isopentane was chilled in liquid nitrogen, ensuring full contact between the two. Brains were embedded in TissueTek within cryomolds, rapidly frozen in the cooled isopentane, transferred to dry ice and stored at -80 °C in a sealed container.

For sectioning, tissues were equilibrated in a -20 °C cryostat and cut into 10-12 µm sections. Each section was mounted onto a Visium gene expression slide by gently touching the active surface, then immediately stored at -80 °C to preserve RNA integrity and spatial organization. Preparations were performed for three mice per condition, with 3-4 brain slices per mouse.

### Visium spatial immunofluorescence staining

Before visium spatial gene expression, the sections were incubated for 1 min at 37 °C and fixed in cold methanol for 30 min at -20 °C. All following steps were carried out in the Visium slide cassette. Sections were blocked with 70 µl of 1x Blocking Buffer (3x SSC Buffer, 2 % BSA, 0.1 % Triton X-100 (w/v), 1 U/µl RNase inhibitor, 10 % Ribonucleoside Vanadyl Complex in H_2_O) for 5 min at RT. After removal, 50 µl of Primary Antibody Solution (3x SSC Buffer, 2 % BSA, 0.1 % Triton X-100, 6.4 U/µl RNase inhibitor in H_2_O with GFP antibody (Aves Lab, GFP-1010) diluted 1:50) was applied and incubated 30 min at RT. Following five washes with 100 µl of Wash Buffer (same composition as Blocking Buffer), sections were incubated with 50 µl of Secondary Antibody Solution (3x SSC Buffer, 2 % BSA, 0.1 % Triton X-100, 6.4 U/µl RNase inhibitor in H_2_O with secondary antibody (diluted 1:100 and DAPI 1:220) for 30 min at RT, followed by another five washes. Slides were removed from the cassette, dipped 20 times in 50 ml of 3x SSC buffer, and covered with 200 µl of Mounting Medium (85 % Glycerol, 2 U/µl Rnase inhibitor in H_2_O) and a coverslip for immediate imaging. Afterwards, coverslips were detached in 3x SSC, and slides were returned to a clean Visium cassette for spatial gene expression analysis.

### Visium Spatial Gene Expression experiment

Following fluorescence staining, spatial transcriptomic profiling was carried out using the “Visium Spatial Gene Expression Slide & Reagent Kit” (10X Genomics) in accordance with the manufacturer’s protocol. Tissue sections underwent controlled permeabilization to release intracellular RNA, followed by reverse transcription and second strand synthesis to generate double-stranded cDNA. After denaturation, cDNA was PCR-amplified and purified to remove by-products. The purified cDNA was then subjected to fragmentation, end repair and A-tailing to create 3’ ends compatible with adapter ligation. Post-ligation cleanup removed excess adapters, and a sample index PCR introduced unique sample indices for multiplexed sequencing.

### Immunohistochemistry imaging of sagittal brain slices

Animals were anesthetized via intraperitoneal injection of 10% ketamine and 2% xylazine, followed by transcardial perfusion with 1x PBS for 8 min and 4% paraformaldehyde (PFA) for 20 min (flow rate 3 ml/min). Brains were dissected and post-fixed in 4% PFA for 30 min at room temperature (RT) followed by incubation in 30% sucrose at 4°C for 2 days. To generate slices, brains were partially embedded in Tissue Tek and frozen at -20 °C for 30 min. Sagittal sections (30 μm) were cut using a Leica CM1950 Cryostat (chamber temperature -20 °C; block temperature -20 °C) and immediately stored in 1x PBS.

Sagittal brain slices were washed three times with 1X PBS for 10 min and blocked/permeabilized with 5% bovine serum albumin (BSA) and 0.5% Triton-X in PBS for 1 h. Primary antibodies were diluted in blocking/permeabilization solution and incubated over night at 4°C. After washing three times with 1X PBS for 10 min secondary antibodies were diluted in blocking/permeabilization solution and incubated for 2 h at RT. Slices were washed three times with 1X PBS for 10 min and incubated in DAPI (dilution 1:1000 in PBS) for 10 min. For quenching slices were dried on a slide and incubated in TrueBlack solution (20X TrueBlack to 1X in 70% ethanol). After washing with 1X PBS, slices were mounted with Fluoromount-G. Imaging was done using a confocal LSM980 microscope and formatted using Imaris 10.1.1. Astrocytes were manually counted and statistical testing was performed via unpaired Student’s t-test in GraphPad Prism 9.5.1.

### Sequencing and alignment of single nucleus samples

Paired 150bp snRNA-seq was performed using the 10X Genomics Gene Expression (GEX) 3’protocol with a NovaSeq 6000 sequencer. For the alignment of reads, a custom reference was created by adding the sequences of the V1S/SV2 transgene and the *Camk2a* promoter to the mm10 mouse reference genome. Count matrices were obtained using the *cellranger count 7.1*(20) pipeline, including introns.

### Sequencing and alignment of visium spatial samples

Sequences were fiducially aligned to spots using Loupe Browser ver. 8. All aligned sequences were mapped using *spaceranger count* 3.0.1 with a custom reference, which included sequences for the promotor and transgene (Camk2aTTA, V1S/SV2) to the mouse genome mm39.

### Quality Control of snRNA

Count matrices generated by *cellranger count 7.1*(*20*) were loaded into an AnnData object and processed using the Python-based framework *Scanpy 1.10.2*(*21*). Integration with R, where needed, was facilitated through the rpy2 package. Raw count matrices were corrected for ambient RNA contamination using the *SoupX 1.6.2*(*22*). To remove potential doublets, *scDblFinder 1.18.0*(*23*) was employed with a fixed seed (123). Nuclei with nUMI and nGenes values exceeding three median absolute deviations (MADs) from the median were excluded. Genes detected in fewer than five nuclei across the dataset were excluded. The resulting dataset was normalized via *scanpy.pp.normalize_total* and *scanpy.pp.log1p*. Highly variable genes were identified using the function *scanpy.pp.highly_variable_genes* with the Seurat v3 flavor, selecting the top 4,000 genes. Dimensionality reduction was performed using principal component analysis (PCA) and batch effects were corrected using the python-implemented version of Harmony via the function *scanpy.external.pp.harmony_integrate*. Harmony embeddings were then used to construct a k-nearest neighbor (kNN) graph with *scanpy.pp.neighbors*. Clustering was performed using Leiden clustering with standard parameters via the function *scanpy.tl.leiden*.

### Quality Control of visium spatial

We filtered each sample of the visium spatial dataset based on the MAD filtering of number of reads (nUMI), number of genes (nGene), and percentage of mitochondrial genes (percent.mt). A spot was filtered out if it was outside of 3x MAD value in at least two metrics. Filtered samples were merged into one Seurat 5.1.0 object(24) and we obtained normalized counts by the *SCTransform* function of Seurat. Integration was performed using *Harmony 1.2.0*(*25*) on 50 PCA embeddings and clustering was done using Leiden clustering based on 30 harmony embeddings. Integrated clusters were visualized using the UMAP method. Samples that were not successfully integrated (based on similarity measures of the harmony embeddings) and showed high percentage.mt or low nUMI levels compared to other samples, were removed from subsequent analysis. A final integration and clustering were performed after filtering.

### Annotation of Cell types in snRNA

Clusters were annotated using literature, the mousebrain (mousebrain.org)(26), and markers identified via the *FindConservedMarkers* function in Seurat. First, neurons and non-neuronal cells were distinguished using mainly canonical markers, such as but not limited to *Rbfox3* (neurons), *Mbp* (oligodendrocytes), *Acsbg1* (astrocytes), *Pdgfra* (oligodendrocyte precursor cells), *Inpp5d* (microglia), *Colec12* (vascular cells), and *Ttr* (choroid plexus cells). Neurons were further classified into Vglut1 (*Slc17a7*), Vglut2 (*Slc17a6*), GABA (*Gad2*), cholinergic (*Scube1*), and dopaminergic (*Th*) neurons. Vglut1 and GABA neurons were further annotated into subtypes based on subclustering and *FindConservedMarkers* markers. A full list of markers and cell types can be found in the Suppl. Table 5.

### Annotation of Regions in visium spatial

Regions were first annotated based on a 0.1 resolution clustering to get high level region annotation (Cortex, Hippocampus, Subcortex). Each high-level region was further annotated based on either more granular resolutions or subclustering. Marker genes from mousebrain (mousebrain.org)(26) and literature were used in combination with the Allen mouse brain atlas(27, 28) to obtain anatomically relevant annotations. All used markers can be seen in Suppl. Table 1.

### Differential Gene Expression Analysis for snRNA

Counts were pseudo-bulked to sample level and filtered for counts above three in at least two samples. Mitochondrial and ribosomal genes were also removed. We created a *Deseq2 1.44.0* object(29, 30) to normalize counts and calculate covariates using *sva 3.52.0*(31). Sva covariates and condition were included into the design (∼ SV1 + condition) to perform Wald testing. P-values were recalculated using *fdrtool 1.2.18*(32) with the “stat” values and adjusted p-values were obtained by Benjamini-Hochberg(33) procedure as applied in the R function *p.adjust*. A gene was considered significantly differentially expressed if the change was below an FDR of 0.05.

### Differential Gene Expression Analysis for visium spatial

Counts were pseudo-bulked to Sample-Region. For the “Spreading Signature” we compared Sample-Spreading Area to Sample-No Spreading Area. Counts were filtered for counts above three in at least three samples and mitochondrial genes were removed. Covariates were calculated using sva on a sample wise paired design, which led to the final design: ∼ SV1 + condition + condition:sample.n + condition:region. Wald testing was performed, p-values were recalculated using fdrtool with the “stat” values and adjusted p-values were obtained by Benjamini-Hochberg procedure as applied in the R function *p.adjust*. Cell number differences between the regions were accounted for by subsampling to a maximum number of cells of two-fold difference. Subsampling was performed accordingly and everything was run 100 times. Only genes that were below 0.05 FDR in at least 95 of 100 runs were deemed as significant. All differential results can be seen in the Suppl. Table 2.

### Ligand-Receptor Analysis for visium spatial

Ligand-Receptor (LR) interactions were calculated using *stlearn 0.4.12*(*34*) on an object of raw counts. Counts were filtered by pp.*filter_genes* function with the *min_cells* parameter of 3. Normalization was performed by pp.*normalize_total* function. The used LR database was downloaded from stlearn via *tl.cci.load_lrs* with species set as “mouse” for the “connectomeDB2020_lit” database. Spatially relevant significant LR pairs were calculated by *tl.cci.run* with parameters *min_spots = 20, distance = None,* and *n_pairs = 10,000* followed by *tl.cci.run_cci* with the parameters *min_spots = 3, spot_mixtures = False, sig_spots = True,* and *n_perms = 1,000*.

### Transcription Factor Analysis

Transcription factors were found using *decoupleR 2.10.0* architecture(35). We used the Deseq2 output for the Spreading Signature as input into the function run_ulm() with the standard parameters. We used the collectTRI mouse database as *net* and set minsize to 5.

### Inference of Networks

Using PIDC, the most interconnected genes were chosen based on T. E. Chan et al.. PIDC, based on Partial Information Decomposition (PID), infers statistical dependencies between triplets of genes from a single-cell gene expression matrix by assigning a confidence score to each interaction. Only the genes with a top 5% confidence score were used to build an interaction network using curated public databases of known interactions. Central genes were defined based on the betweenness centrality scores calculated via the *igraph 2.0.3*(*36*) architecture.

### Finding Interactions in interaction databases

The OmniPath database(37) was accessed using *OmnipathR 3.11.16*(*37*) and combined with interactions having a confidence score above 0.600 from full network type STRING(38). We included protein complexes and post translational modifications from OmniPath as interactions. An interaction was considered “Found” in the database, if the connection had a maximum of one bridge gene between the two interactors.

### Spot deconvolution

Visium spatial spots were deconvoluted using an annotated snRNA dataset from the same mouse model. We used the *CARD 1.1* package(39) to deconvolute sample wise each spot for astrocyte, microglia, oligodendrocyte, OPC, vascular cells, choroid plexus, cholinergic neurons, dopaminergic neuron, GABAergic neuron, Vglut1 neuron, and Vglut2 neuron. The default parameters were used for creating and deconvoluting CARD object with minCountGene set to 100 and minCountSpot to 5.

### Analysis of Cell type composition differences

Cell type composition was based on the percentages from deconvoluted spots by CARD. Most probable changes were calculated using *tascCODA 0.1.3*(*40*) by using all cell types as a reference once. Changes were deemed significant if it was changed in at least 66% of reference iterations with a maximum p-value of 0.2.

### Module Score Analysis

Module score of “Spreading Signature” for snRNA dataset was obtained using the Seurat function AddModuleScore. The difference of Module Scores between conditions per cell type was tested for significance using the Wilcoxon-mann-Whitney test(41) via the R function *wilcox.test().* We calculated the effect size for each comparison using the *cliff.delta()* R function(42).

### Term Enrichment analysis

GO and KEGG term enrichment was performed using the R packages *clusterProfiler 4.12.1*(*43*) and *pathfindR 2.4.1*(*44*) respectively. Both were run with default parameters, with minGSSize for enrichGO function set to 15. Enriched KEGG terms containing one of the terms “Addiction, Corona, Infection, Invasion, Cancer, Alcohol, Chagas disease, Cushing, Glioma, Melano, Oocyte, Hepatitis, Tuberculosis, African, Viral, Carcinoma, Circadian, Cardiomyocytes, Leukemia, cardiomyopathy” were not considered for downstream analysis. For enriched GO or KEGG terms from Vglut1 subtypes we only considered terms relevant containing one of the terms “synapse, synaptic, neurotransmitter, vesicle, transmitter, astrocyte, astrocytic, transmembrane signaling, cell adhesion”

### Comparing with Human data

We used three published human snRNA-seq datasets(16, 17, 45). For comparing mouse Astro-Slc1a2 data to human data, we used the original annotation from the original publications, isolated the Astrocyte, ran *Deseq2 1.44.0*(29, 30) on each dataset (pseudo-bulked on sample level) and compared the log2 fold changes to the mouse data. For comparing the “Spreading Signature” we pseudo-bulked every human dataset on a sample level and ran *Deseq2 1.44.0*(29, 30) to compare log2 fold changes.

## Results

### Defining a Spreading Area and “Spreading Signature”

To study the spatial characteristics of α-syn spreading in PD, we utilized our transgenic α-syn PD mouse model that recapitulates α-syn oligomer propagation and accumulation at synapses within specific forebrain and midbrain regions, including the SN(18).

Transcriptional changes in response to α-syn oligomer spreading were determined using visium spatial transcriptomics on brain slices of 20-months (20M) and 3-months (3M) old mice from the previously described PD mouse model with an age and α-syn oligomer dependent motoric phenotype (Additional File 1), which is based on a human α-syn protein complementation system expressing α-syn fused to non-fluorescent Venus YFP halves (V1S and SV2). Fluorescent Venus YFP (V1S/SV2) is reconstituted when α-syn oligomerization takes place (Figure 1 a). We included two conditions: Double-transgenic mice expressing V1S/SV2 under the control of the tTA-Camk2a transgene (ON) and single-transgenic mice lacking the tTA-Camk2a transgene (OFF) as control (Figure 1 a). Our model enabled the visualization of the presence of α-syn protein (V1S/SV2 signal) on a spatial level (Figure 1 b) and visium spatial transcriptomics supplying the corresponding mRNA levels of V1S/SV2 (Figure 1 c).

**Figure 1.**
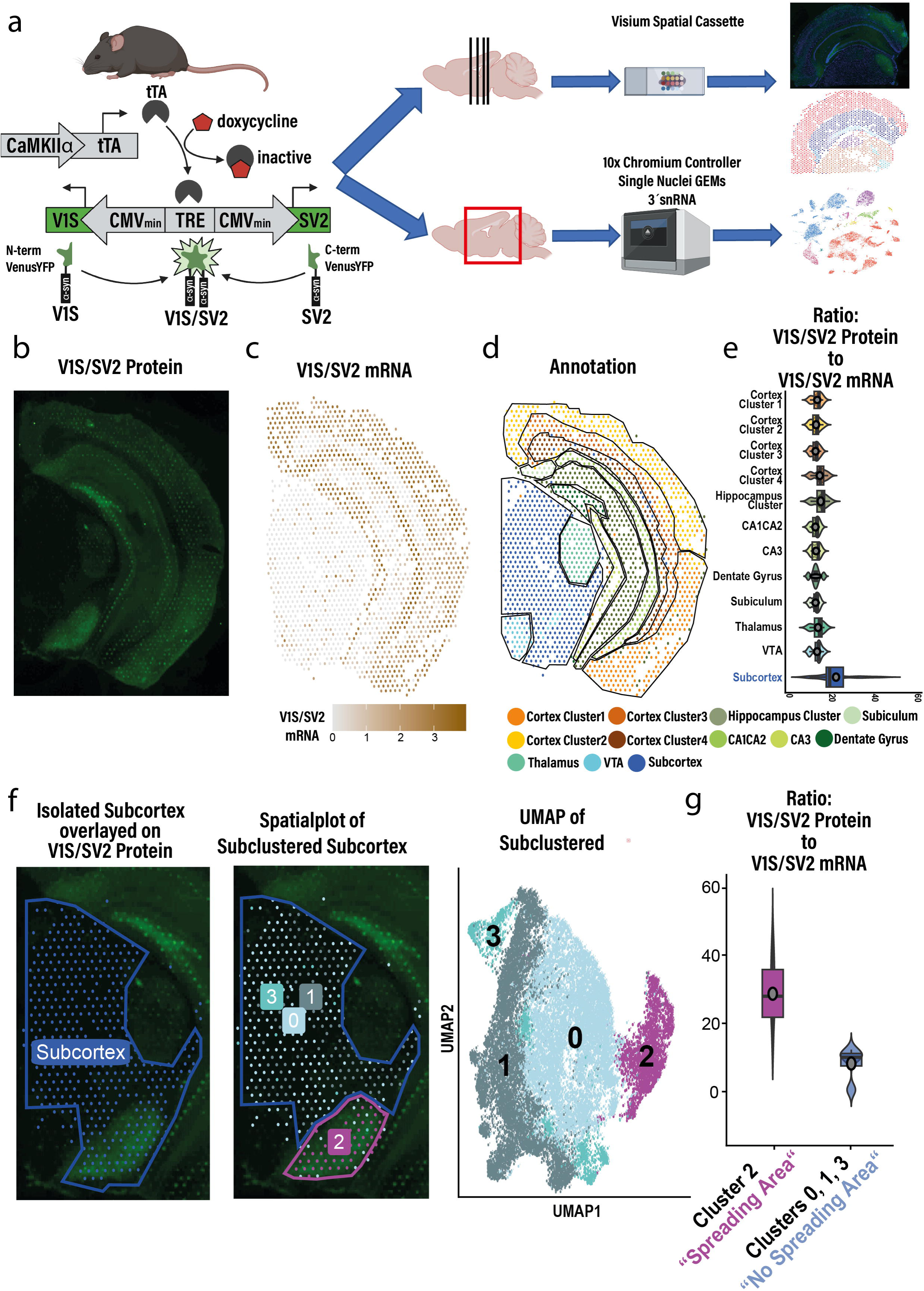
Identification of a “Spreading Area” with spreadingα-syn oligomers in an *in-vivo* PD mouse model. **a,** Schematic representation of the experimental design. Brain tissues from aged mice in a Venus-tagged human α-syn oligomer-based PD model was used to generate two datasets: Visium spatial transcriptomics and snRNA. Both datasets included two conditions: ON (double-transgenic mice with V1S/SV2 α-syn oligomer expression under the control of tTA-Camk2a transgene) and OFF (single-transgenic mice without tTA-Camk2a). The visium spatial dataset comprised 3–4 brain slices per mouse, with samples from three mice per condition, while the snRNA-seq dataset included samples from three mice per condition. **b,** Using the V1S/SV2 signal, which only emerges if at least two α-syn combine together, we evaluated visually the presence of V1S/SV2 proteins on mouse brain slices. c, The mRNA levels of V1S/SV2 was also obtained on a spatial level using visium spatial transcriptomics. d, Brain slices in the visium spatial dataset were annotated using Louvain clustering, regional markers obtained from mousebrain (mousebrain.org)(26) or comparing obtained markers from Wilcoxon rank sum test to published markers (Additional File 3), and anatomical information from the allen mouse brain atlas(27, 28). This approach enabled the assignment of anatomically relevant cluster annotations. **e,** As we used the same brain slices for V1S/SV2 Signal and V1S/SV2 mRNA levels, we could calculate a protein to mRNA ratio per annotated region. The Subcortex showed the highest protein to mRNA ratio, suggesting V1S/SV2 spreading into the Subcortex. f, To further characterize the Subcortex we subclustered the Subcortex and obtained four clusters. Notably, Cluster 2 was the sole cluster to be regionally distinct and representative of the region with the highest V1S/SV2 protein levels. g, Thus, we split the annotation of the Subcortex into the regions of interest (ROI) “Spreading Area” and “No Spreading Area”. Indeed, the higher ratio between signal and mRNA was clearly driven by the “Spreading Area”.

After filtering and clustering the integrated data (Additional File 1) we annotated spatially distinct cell clusters based on transcriptional mouse brain markers(26) and regional annotations from the Allen mouse brain atlas(27, 28) (Figure 1 d, Additional File 2 a, Additional File 3).

As we used the same brain slices for both V1S/SV2 protein and V1S/SV2 mRNA detection, we calculated the ratio between the V1S/SV2 protein (mean Venus intensity by ImageJ(46)) and V1S/SV2 mRNA levels (Figure 1 e). Notably, the Subcortex showed elevated ratio levels, implying α-syn oligomer spreading to the Subcortex (Figure 1 e).

To further investigate the Subcortex we subclustered the Subcortex into 4 transcriptionally distinct clusters (Figure 1 f). Only cluster 2 showed a spatially distinct character and was representative of the region with a high ratio between V1S/SV2 protein to V1S/SV2 mRNA levels (Figure 1 f, g). We named the cluster 2 “Spreading Area” and the rest of the Subcortex “No Spreading Area” (Figure 1 g). It should be mentioned that the “Spreading Area” overlapped with both the substantia nigra (SN) pars compacta and pars reticulata and non-SN regions based on Allen mouse brain atlas(27, 28) (Additional File 2).

This distinction provided a framework to investigate transcriptional changes associated with the spreading of α-syn oligomers. By comparing the transcriptional profiles of the “Spreading Area” and “No Spreading Area” we aimed to elucidate transcriptional consequences of propagation of α-syn oligomers.

For the characterization of the “Spreading Area”, we defined a “Spreading Signature” comprising two components: First, differentially expressed genes (DEGs) were identified by comparing the “Spreading Area” and “No Spreading Area” regions (Additional File 4, Additional File 5). Only DEGs that were significant in the ON condition but not in the OFF condition were included (Figure 2 a, Additional File 6). Second, using the aforementioned DEGs, transcription factors (TFs) that actively regulate these DEGs were inferred (Figure 2 a). Together, these analyses resulted in a transcriptional “Spreading Signature” encompassing a total of 135 genes/TFs (Figure 2 a, Additional File 7).

**Figure 2.**
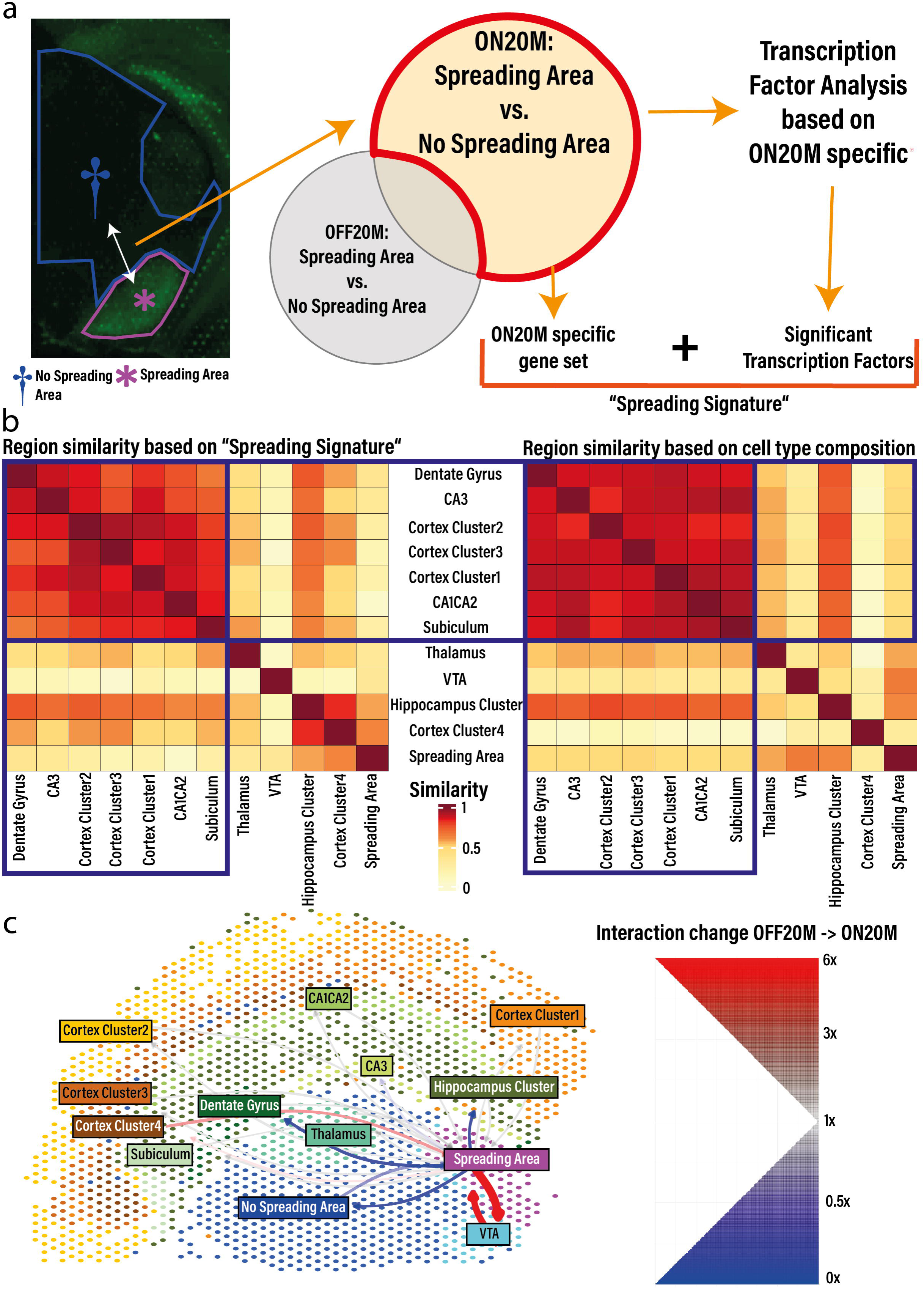
α-syn oligomer spreading related transcriptional changes differ between “Spreading Area” and α-syn mRNA expressing regions. **a,** To investigate transcriptional changes associated with spreading, we conducted a differential gene expression analysis comparing the “Spreading Area” and “No Spreading Area” regions. We retained only DEGs that were unique to the ON condition (”ON-Spreading Area” vs. “ON-No Spreading Area”), subtracting those already identified in the OFF condition (”OFF-Spreading Area” vs. “OFF-No Spreading Area”). These ON-specific DEGs were subsequently analyzed for TF activity using the decoupleR(35) framework. Relevant TFs and the 20M ON-specific DEGs were consolidated to define the “Spreading Signature.” **b,** Heatmaps depicting similarities between annotated regions are shown. The left heatmap illustrates region similarities based on Pearson correlation of the log2 fold changes between annotated regions and “No Spreading Area” of the “Spreading Signature” genes. Notably, the “Spreading Signature” was absent in regions with native high expression of V1S/SV2 mRNA. This grouping pattern was mirrored in the right similarity heatmap for the high V1S/SV2 mRNA level regions, derived from the Aitchison similarity of the deconvoluted cell type compositions of every annotated region. This grouping pattern suggests that transcriptional differences between regions are partially explainable by variations in cell type composition for cortical and hippocampal regions. **c,** Changes in region-to-region LR interactions from OFF to ON conditions visualized on an exemplary sample. Notably, interactions between “Spreading Area” and the ventral tegmental area (VTA) increased nearly six-fold.

To examine whether the “Spreading Signature” was unique to the “Spreading Area” we performed likeness analysis by comparing how the “Spreading Signature” was expressed in every annotated region compared to the “No Spreading Area”. We found that the “Spreading Signature” was similarly changed in the “Spreading Area”, “Cortex Cluster 4”, and “Hippocampus Cluster” (Figure 2 b heatmap left). The other cortical and hippocampal regions with high mRNA levels of V1S/SV2 did not show similarities to the “Spreading Area” but instead obtained high correlation between each other (Figure 2 b heatmap left). This indicates that the α-syn oligomer spreading related transcriptional changes in the “Spreading Area” are not present in regions with high mRNA expression of V1S/SV2.

We hypothesized that these differences were primarily attributable to variations in cell type composition between regions, thus visium spatial spots were deconvoluted using snRNA-seq data from the same mouse model and cell type compositions between the regions were compared. Indeed, the comparison indicated that the differences in the “Spreading Signature” between the “Spreading Area” and the cortical or hippocampal regions were largely explained by cell type composition differences (Figure 2 b heatmap right, Additional File 6).

Additionally, ligand-receptor (LR) analysis revealed significant alterations in interregional communications between different regions in response to V1S/SV2 oligomers mainly involving the “Spreading Area” (Figure 2 c, Additional File 8). Notably, intercellular communication between the “Spreading Area” and the tyrosine hydroxylase (TH) enriched ventral tegmental area (VTA) increased nearly six-fold in 20M mice when α-syn oligomers were present, pointing to a strong impact of α-syn oligomers on the interactions between VTA and the “Spreading Area” (Figure 2 c).

### **α**-syn spreading promotes astrocyte/neuron dysfunction

To understand the biological and molecular meaning of the “Spreading Signature”, we constructed a transcriptional network with PIDC(47) using the “Spreading Signature” and transcriptional data from the “Spreading Area” in 20M ON mice. We found that the resulting network was composed of two subnetworks (Figure 3 a): The first subnetwork was predominantly associated with astrocytic uptake and neurotransmitter processing, including genes such as *Slc1a2*, an astrocyte-enriched glutamate/excitatory amino acid transporter (EAAT2) critical for glutamate clearance and processing at synapses (Figure 3 a, Additional File 7). The second subnetwork was linked to astrocyte–neuron interaction dysfunctions, such as the astrocyte-neuron pre-protein/neurotransmitter *Penk* (Additional File 7). Together, network analysis suggested a central role of astrocytes in α-syn spreading.

**Figure 3.**
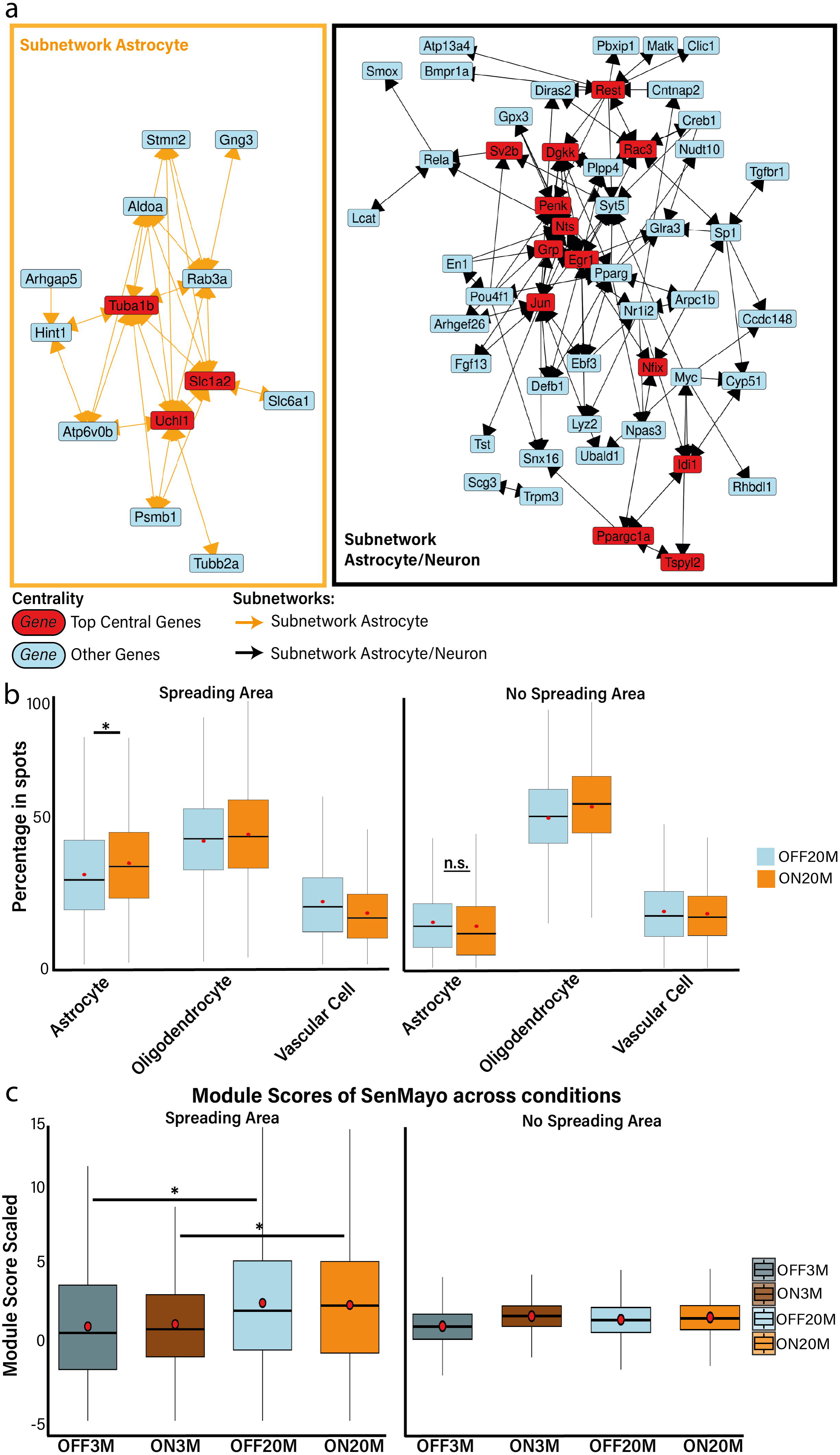
*In-silico* analysis indicate central role of astrocytes in response to α-syn oligomer spreading. **a,** A network was constructed using the “Spreading Signature,” transcriptional data from the “Spreading Area”, and known interactions from omniPath(37) and STRING(38) via the PIDC(47) algorithm. The network revealed two distinct subnetworks, with central genes identified based on betweenness centrality scores for both subnetworks. Functional annotations of the network genes, linked both subnetworks to astrocyte–neuron interactions, including astrocytic uptake and neurotransmitter glutamate processing (Suppl. Table 3). **b,** Significant changes in cell type composition between the OFF and ON conditions were assessed by taking the deconvoluted cell type ratios per region as cell type numbers and using tascCODA(40). Shown here are the top three most represented cell types in the “Spreading Area”. Astrocytes exhibited a significant (*: FDR < 0.2 in 60% of tascCODA(40) iterations) increase in numbers in the “Spreading Area”, whereas no significant changes were observed in the “No Spreading Area”. **c,** Comparison of senescence levels across conditions and regions using the SenMayo(48) gene set. Scores are scaled to the mean of 3M OFF set to one. We only observed in the “Spreading Area” a significant (unpaired t-test on a sample level, * FDR < 0.05) increase in senescence between the age groups. “No Spreading Area” showed no significant changes.

To further elucidate the central role of astrocytes, we investigated whether the astrocyte numbers were altered in the “Spreading Area” between On and OFF conditions. We used the deconvoluted ratios of cell types per region (Additional File 9) as cell type numbers to run tascCODA(40) between conditions. Indeed, we could observe a significant increase of astrocyte numbers in the “Spreading Area” in response to α-syn oligomers (Figure 3 b). This increase in astrocyte numbers was not observed in the “No Spreading Area”.

Furthermore, we also found senescence changes to differ between the “Spreading Area” and “No Spreading Area”. We employed the SenMayo gene set(48) and studied possible changes of senescence in the “Spreading Area” between ages and α-syn oligomer presence. We observed a significantly higher enrichment of the SenMayo gene set in the “Spreading Area” during aging (Figure 3 c), while α-syn oligomer expression had no notable effect on senescence (Figure 3 c). The “No Spreading Area” showed no significant changes in the senescence during aging nor as a response to α-syn oligomer expression (Figure 3 c). This suggests that the “Spreading Area” has a predisposition being vulnerable for age related changes compared to the “No Spreading Area”. This might be responsible for the accumulation of spread α-syn oligomers in the “Spreading Area” eventually leading to the observed transcriptional changes in astrocytic function (Figure 3 a).

To further characterize interactions between astrocytes and α-syn oligomers in the SN and to verify the increase of astrocyte numbers, we conducted immunohistochemical (IHC) staining of 20M brain slices of our V1S/SV2 mice visualizing astrocytes, α-syn oligomers, and dopaminergic neurons (Figure 4, Additional File 10). Manual counting indeed confirmed our *in-silico* findings and revealed a significant increase in astrocyte numbers in response to α-syn oligomers in the SN of aged PD mice. Moreover, astrocytes displayed morphological changes towards hypertrophic astrocytes with thickened and elongated processes. (Figure 4 a, b).

**Figure 4.**
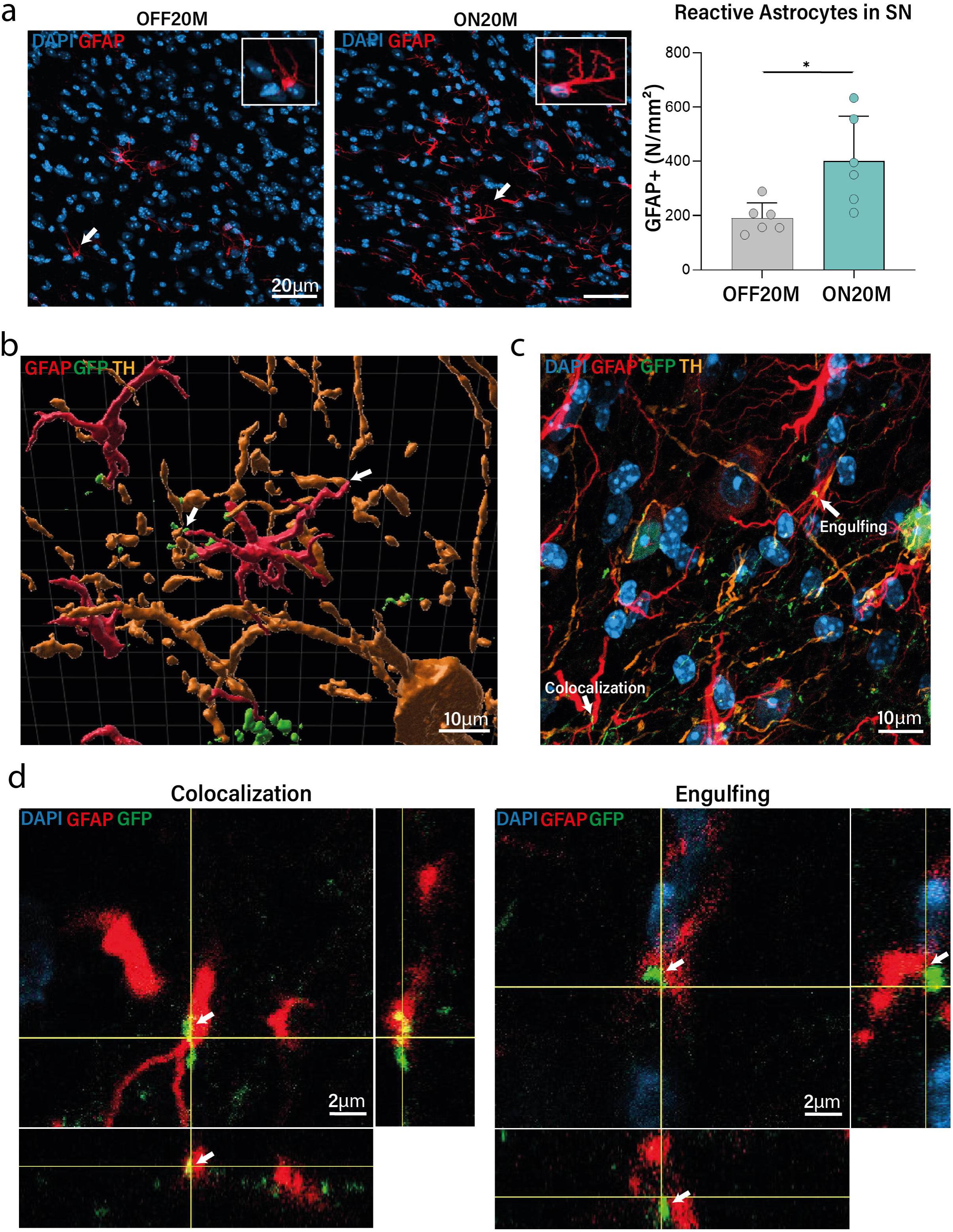
Expression of α-syn oligomers leads to increase in astrocytes in the SN. **a,** Using brain slices from 20M mice from our PD model we performed IHC staining of astrocytes (GFAP, red) and cell nuclei (DAPI, blue). Number of astrocytes were manually counted. Microscopy images on the left show representative images of the OFF and ON conditions with white arrows highlighting astrocytes included in the count. White boxes are zoomed in images of the white arrow astrocytes. Quantification of astrocytes in the SN revealed a significant increase in their numbers in response to α-syn oligomer expression (n = 6 per condition; mean ± SD; unpaired t-test, *p < 0.05, 40× magnification). **b,** Surface rendering of astrocytes, V1S/SV2 α-syn oligomers (Venus signal amplified by GFP, green), and dopaminergic neurons (TH, orange) staining using the Imaris software on a representative 20M ON brain slice. We observed contacts and close proximity between neurons, astrocytes and V1S/SV2. **c,** Z-stack maximum intensity projections of IHC staining in 20M ON SN showed two distinct localization patterns of the V1S/SV2: (Engulfing) V1S/SV2 was found in close proximity to astrocytes being enclosed but not taken up by folding astrocytes and (Colocalization) colocalized within astrocytes. **d,** Magnified orthogonal projections of “Colocalization” and “Engulfing” arrows in C. Bottom image: XZ-plane, right image: YZ-plane, middle image: XY-plane. DAPI: Nucleus, GFAP: astrocytes, GFP: V1S/SV2, TH: Dopaminergic neuron

Extracellular α-syn was mostly found at the end feet of astrocytes and synaptic ends (Figure 4 b). We further observed two distinct localization patterns of α-syn oligomers in the context of astrocytes: (1) Colocalization: α-syn taken up by astrocytes and (2) Engulfing: α-syn engulfed by astrocytes but not taken up (Figure 4 c, d).

While visium spatial offers reliable spatial transcriptomic information, it does not offer cell type specific data. Thus, to further strengthen the central role of astrocytes affected by α-syn spreading we utilized snRNA-seq data from the same mouse model at 23 months to obtain cell type specific data. After thorough quality control and annotation (Additional File 11, Additional File 12), we quantified the expression of the “Spreading Signature” (split into up and down regulated) across all cell types and conditions in our snRNA-seq dataset (Figure 5 a). Only the astrocytic subtypes - Astro-Slc1a2 and Astro-Slc6a1 - exceeded the significance threshold (FDR < 0.05 & Effect size > 0.15; Figure 5 a) but notably only for the up regulated “Spreading Signature” genes. This again cemented the strong role of astrocytes in α-syn spreading found by visium spatial data analysis with an added focus on the up regulated genes in the context of astrocytes.

**Figure 5.**
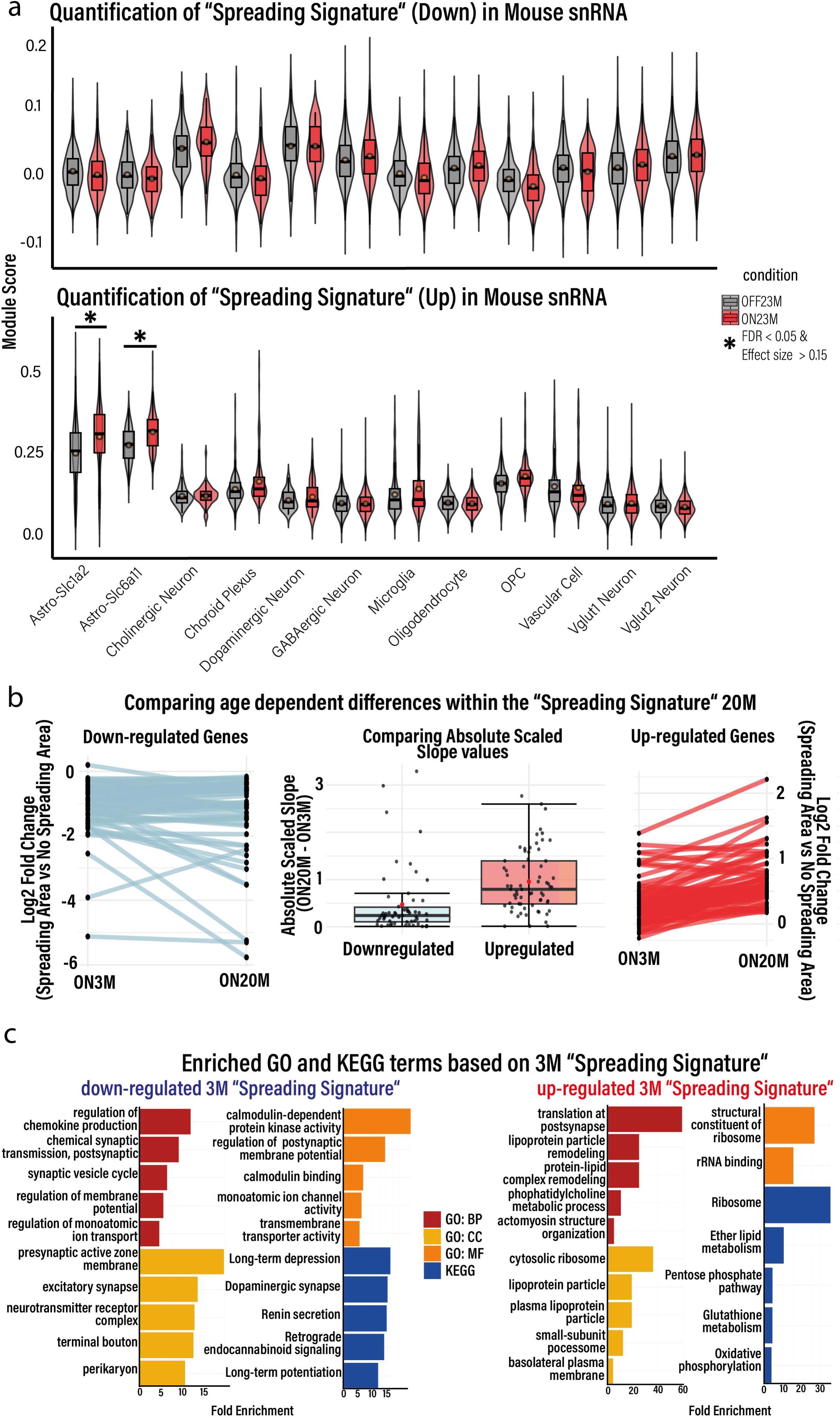
Motoric deficits and α-syn oligomer spreading related transcriptional changes of astrocytes are age related. **a,** Using snRNA data from 23M old mice of our PD mouse model we calculated a Module Score of our “Spreading Signature” for all annotated cell types using the Seurat(24) function *AddModuleScore*. Comparing the scores between the conditions, only the astrocytic subtypes cleared the significance threshold of FDR < 0.05 and effect size (cliff delta) > 0.15 (Wilcoxon rank sum test) in the up regulated “Spreading Signature” genes. **b,** We investigated our “Spreading Signature” of interest from 20M across the ages by comparing the log2 Fold Changes of the “Spreading Area” vs. “No Spreading Area” comparison in the visium spatial data between 3M and 20M. While the down-regulated genes were similar in the 3M and 20M, we observed heightened up-regulation in the up-regulated genes at 20M. **c,** We checked the function of the 3M “Spreading Signature” by GO and KEGG term enrichment analysis. For visualization enriched GO and KEGG terms (FDR < 0.05) were clustered by similarity and only representative terms of the clusters were visualized. Down-regulated genes were linked to neuronal, synaptic, and calcium signaling terms while up-regulated genes were enriched in ribosomal, cytoskeleton, and mitochondrial functions

Taken together, using spatial transcriptomics data, IHC data, as well as snRNA-seq data we could show the increase of astrocyte numbers, their interaction with α-syn oligomer in the SN in aged PD mice, and the importance of astrocytes in the context of α-syn oligomer spreading.

### Astrocytic dysfunction driven by **α**-syn spreading is correlated with aging

Next, we went back to the visium spatial data to analyze the “Spreading Area” also in younger mice (3M) and asked whether related transcriptional changes also occur in young mice. Using the same methods as for the 20M PD mice, we confirmed α-syn oligomer spreading in the same region in the 3M PD visium spatial data (Additional File 14). Notably, the down-regulated genes from the 20M “Spreading Signature” were similarly regulated in 3M and 20M while, the up-regulated genes were highly elevated upon aging (Figure 5 b). We had already shown that especially the up regulated genes in the “Spreading Signature” from the 20M old animals are enriched in the astrocyte subtypes (Figure 5 a). This suggests that aging amplifies the transcriptional changes of astrocytes in response to α-syn oligomer spreading.

To get a more holistic transcriptional comparison between the age groups, we determined the transcriptional characteristics of the 3M “Spreading Area” and defined a “Spreading Signature” for the 3M old mice as for the 20M mice (Additional File 16, Additional File 17). There was only a miniscule overlap between the 20M “Spreading Signature” and the 3M “Spreading Signature” (Additional File 15).

We further investigated the function of the 3M “Spreading Signature” and found genes related to neuronal, synaptic, and calcium signaling function to be down-regulated, while up-regulated genes were involved in ribosome, cytoskeleton, and mitochondrial function (Figure 5 c).

Taken together, our findings suggest that the neuronal function has some disruptions already at 3M that further progress during aging (Figure 5 c). During aging, astrocytes get affected by α-syn oligomers (Figure 4; Figure 5 a, b; Additional File 15), which may enhance the disruptive load on neuronal function (Figure 5 c).

### Dysregulation of the astrocyte-neuron glutamate system

As the Astro-Slc1a2 subtype had most of the significant DEGs when comparing ON vs OFF conditions in astrocytes (Additional File 13) and due to the presence of *Slc1a2* as a central gene in the “Spreading Signature” (Figure 3 a) we focused further downstream analysis on the Astro-Slc1a2 cell type.

Our special interest was to study a potential impact of astrocyte dysfunction on neurons. We used our snRNA-seq data from 23M old mice and obtained DEGs between PD ON vs. OFF. Using the DEGs we performed Gene Ontology (GO) and KEGG pathway enrichment analyses.

GO and KEGG pathway analysis highlighted glutamatergic synapses, neurotransmitter cycles and TCA-cycle metabolism (Figure 6 a). Furthermore, we constructed a network for the Astro-Slc1a2 DEGs from the mouse snRNA-seq data (Additional File 15) and found *Grm5* as a prominent central gene which corresponds to the metabotropic glutamatergic receptor 5 (mGluR5) on a protein level that plays an essential role in glutamate signaling and astrocyte-neuron interactions(49–51) (Additional File 18).

**Figure 6.**
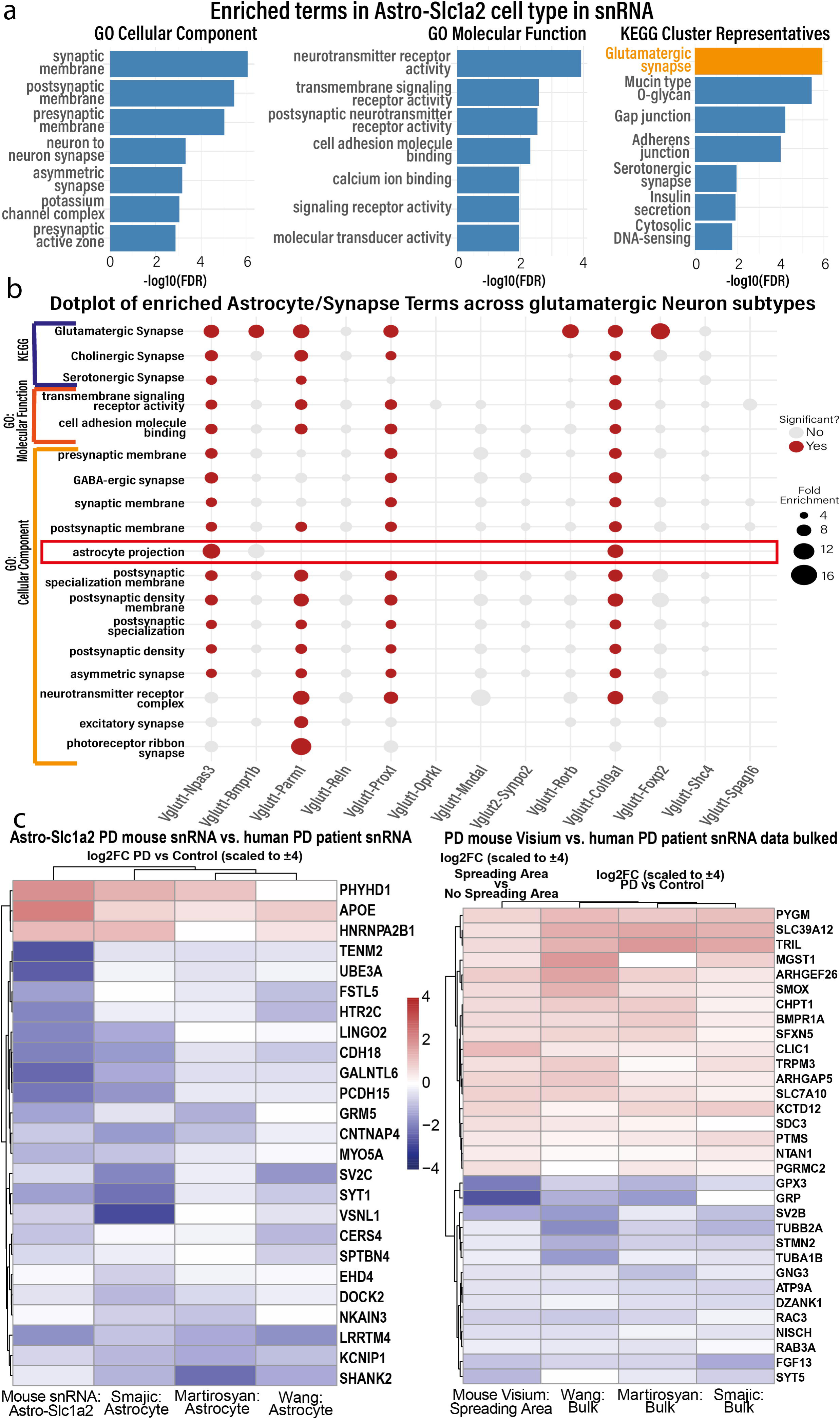
snRNA shows α-syn oligomer related disruptions in the Astrocyte-glutamatergic neuron axis (Astro-GluN) **a,** Enriched Gene Ontology (GO) and KEGG terms of the Astro-Slc1a2 DEGs showed terms related to glutamatergic synapses, neurotransmitter cycles and Astro-GluN axis. **b,** GO and KEGG pathway enrichment analyses were performed on DEGs from each glutamatergic subtype. We filtered for significant terms (padj < 0.05) and visualized particularly terms related to “synapse” or “astrocyte” (s. Methods: *Term Enrichment analysis*). Notably, only the Vglut1-Npas3 and Vglut1-Col19a1 subtypes included “astrocyte projection” among its significant terms, highlighting a unique link between these subtypes and astrocytic processes. **c,** Comparison of log2 foldchanges of Astro-Slc1a2 DEGs between mouse and human snRNA datasets. We employed three published human snRNA-seq datasets(16, 17, 45) from the substantia nigra of PD patients and age-/sex-matched controls. Left compares the log2 fold changes of Astro-Slc1a2 DEGs from our snRNA-seq data to the log2 fold changes between PD and Control in astrocytes in each human dataset. The right compares the log2 fold changes between “Spreading Area” and “No Spreading Area” at 20M ON from our mouse visium spatial data to the log2 fold changes between PD and Control in each bulked human dataset. We saw 25 genes that were changed concordantly on the left in all datasets and 32 genes on the right.

Together, we found *Grm5* – essential for glutamate signaling – as a central gene in the vulnerable *Slc1a2* enriched astrocyte subtype, indicating dysfunctions in the astrocyte-glutamatergic neurons (Astro-GluN) interactions in response to α-syn oligomers.

Focusing on glutamatergic neurons, we used our mouse snRNA-seq dataset and annotated 13 distinct glutamatergic neuron subtypes (Additional File 11) and studied GO and KEGG terms for each glutamatergic subtype. Interestingly, we obtained the term “astrocyte projection” as a significantly enriched term in the subtypes Vglut1-Npas3 and Vglut1-Col19a1 (Figure 6 b). Both cell types also significantly downregulated *Grm3* (Additional File 19, Additional File 20), a gene transcribing the metabotropic glutamate receptor 3 (mGluR3). This aligned with our previous observations since astrocytes already pointed to mGluR5 to be central in α-syn oligomer related changes. These findings suggest that those two subtypes are more affected by astrocytic changes and potentially to the aforementioned dysfunctions in the Astro-GluN axis, with mGluRs possibly playing a pivotal role in the transmission of pathology between astrocytes and neurons (Additional File 21, Additional File 22).

Based on our findings in PD mice including analysis of visium spatial data, snRNA-seq data, and confocal imaging, we could center the effects of α-syn oligomer spreading to astrocytes and changes in the Astro-GluN axis in two distinct glutamatergic subtypes.

### Astro-GluN concordant changes between mouse and human data

In the next approach we set out to test our “Spreading Signature” and Astro-Slc1a2 DEGs in human data from PD patients.

We employed three published human snRNA-seq datasets (16, 17, 45) from the substantia nigra of PD patients and age-/sex-matched controls. First, we compared the log2 fold changes of Astro-Slc1a2 DEGs from our mouse snRNA-seq data to the log2 fold changes between PD and Controls in astrocytes in each human dataset.

While the DEGs exhibited moderate concordance overall, we obtained 25 genes that were changed concordantly in all datasets (Figure 6 c). This included central genes from our snRNA-seq mouse network (Additional File 15), particularly those involved in the Astro-GluN axis such as *GRM5, CNTNAP4,* or *SV2C* (Figure 6 c left heat map).

To compare our “Spreading Signature” from 20M visium spatial PD mice data, we pseudo-bulked the human datasets(16, 17, 45) to a sample level. This was done to make the cell type unspecific “Spreading Area” of our visium spatial data more comparable to the cell type specific snRNA human data. We then compared the log2 fold changes between “Spreading Area” and “No Spreading Area” at 20M from our mouse visium spatial data to the log2 fold changes between PD and Control in each pseudo-bulked human dataset.

General concordance was again moderate, but we saw 32 genes changed concordantly in all datasets (Figure 6 c). With genes for neuronal and synaptic functions (*SV2B, RAB3A, SYT5, RAC3, GNG3, TUBB2A, TUBA1B)* being downregulated and genes related to glial activation (*TRPM3*, *BMPR1A*, *SDC3*) being upregulated.

These results suggest that the Astro-GluN axis, including the aforementioned *Grm5*, at least partially is affected in human PD patients.

## Discussion

Based on multiple modalities this study demonstrates transcriptional changes of the SN in response to α-syn oligomer spreading. We show importance of astrocytes, relevant transcriptional changes in astrocytes related to the Astro-GluN axis, and affected subtypes of glutamatergic neurons. Notably, we observed that these changes in astrocyte changes mainly occur in aged PD mice. Based on our findings, we propose a model of α-syn oligomer spreading affecting the astrocyte GluN axis and report relevant genes in this network (Additional File 22).

Multiple studies consistently reported the importance of astrocytes in PD and their interactions with α-syn(52–58). Moreover, Loria et al.(55) were able to demonstrate a role for astrocytes in degradation of pathological α-syn fibrils in co-cultures of primary neurons and astrocytes. In line with this, EVs derived from erythrocytes and containing α-syn oligomers were shown to accumulate in astrocyte endfeet impairing glutamate uptake likely via EAAT2 channel when injected into mice(59).

Extending on previous findings, we show that in response to α-syn oligomer spreading, astrocytes increase in numbers and change their transcriptional profile in the SN. We report a gene program in the SN specifically affected by α-syn oligomer spreading in which its molecular architecture could confirm the previously reported astrocytic changes involving *Slc1a2*/EAAT2 (60) and introduce new central genes in the α-syn oligomer spreading specific transcriptional profile changes.

Furthermore, previous studies already reported the ability of α-syn oligomers to affect astrocytic functions in recycling glutamate(54, 61, 62), such as Trudler et al.(62) showing that α-syn oligomers induced astrocytes to release more glutamate and thereby increase glutamatergic signaling in human primary neurons from iPSCs. Our data complement these studies by downregulation of genes related to glutamate receptors, astrocytic junctions, and glutamatergic signaling in both astrocytes and affected glutamatergic neurons. This might offer specific central targets to rescue the Astro-GluN axis. Indeed, we point to mGluRs to play a major role in the transition of damage from astrocytes to GluNs and in the overall disruption of the Astro-GluN axis, as we found mGluRs (*Grm3* and *Grm5*) as central players in either the affected astrocytes or the glutamatergic neuron subtypes. Importantly, mGluR3 and mGluR5 have already been shown to be of importance in human astrocytes for the Astro-GluN axis(63).

Previously, mGluRs have been implicated in PD multiple times(64–66) with for example mGluR5 antagonists being protective of dopaminergic neurons in PD animal models(67, 68). We align mGluRs into α-syn oligomer spreading related Astro-GluN dysfunctions and speculate that the disruptions in mGluR signaling might affect recycling of glutamate via astrocyte, and blockades due to accumulation of α-syn oligomers in astrocytes, neurons, and extracellular space, lead to diminished glutamatergic signaling between the synapses. Following the synapse formation/elimination rules(69), the synapses and neurons with less signaling activity devolve and are eliminated.

We further suggest that the involvement of astrocytes in our model only occur with sufficient aging. We observed higher vulnerability to age related effects in our “Spreading Area” by enrichment of the SenMayo gene set pointing to senescent alterations. Whether and how senescence-driven mechanisms contribute to α-syn accumulation and Astro-GluN dysfunctions has to be determined in future studies. It might be that senescent glial cells may facilitate the cell-to-cell spread of pathological proteins and senescence is surely well established as a factor in aging(70) and age related neurodegenerative diseases like PD or Alzheimeŕs Disease(71, 72). Of note, α-syn aggregates have been shown to spread in patterns correlating with regional senescence markers in PD(73). Thus, our data further emphasize astrocytes as an age dependent player in the development of PD phenotypes through α-syn oligomer spreading. Our findings are supported by previous studies that showed reactive astrocyte/astrocytic vulnerabilities are highly correlated with aging in mouse and human brains(74–76).

Here we report a unique gene set found to be central players in the transcriptional changes of astrocyte-GluN network in response to α-syn oligomer spreading. These include down-regulated genes related to glutamatergic signaling, astrocyte-neuron junctions, and protein transport system and increased expression of genes involved in activation of astrocytes, phagocytosis, synapse elimination, and elevated oxidative stress.

In summary, we show the importance of the Astro-GluN axis dysfunction in the context of spreading of α-syn oligomers and provide central genes (Additional File 22) in spreading processes involved for further research to understand PD related Astro-GluN dysfunction.

### Conclusions

In this study we investigated the transcriptional changes to α-syn oligomer spreading in the SN using spatial transcriptomics and snRNA-seq with a PD mouse model. We identified a transcriptionally distinct “Spreading Area” with age- and α-syn related astrocytic changes. We further describe the transcriptional changes to affect the Astro-GluN axis with glutamatergic subtypes being affected by astrocytic transcriptional disruptions and show that some Astro-GluN genes are changed concordantly in both mouse and human SN snRNA-seq datasets. Based on our findings we propose a model of Astro-GluN axis changes due to α-syn oligomer spreading with central genes, such as *Grm3* and *Grm5*, to investigate further.

## Limitations of the study

While our findings are supported by multiple modalities, they are based on our PD mouse model and even though our comparison with human datasets gives an indication of translational effect of our findings in human PD patients, a study with more PD patients is necessary to truly validate the findings. Indeed, the comparison between mice visium spatial data to the bulked snRNA-seq human data has multiple limitations. First, it is not given that a bulked snRNA-seq data is representative of the cell type composition on a visium spatial slide. Second, we compare the log2 fold changes in visium spatial between regions to the log2 fold changes in human snRNA-seq data between conditions. Even though, both comparisons compare “PD pathology to non-PD pathology” the pathology difference between conditions and the pathology difference between regions might differ. Moreover, a future study might include a more granular spatial transcriptomics method to enable true single cell resolution spatial transcriptomics.

## Availability of data and materials

The raw sequencing data that was generated in this study is available from ArrayExpress with the identifier <u>E-MTAB-15216</u> for visium spatial data and the identifier <u>E-MTAB-15158</u> for snRNA data. Detailed metadata, processed seurat objects, counts, spatial images and so on can be found on Zenodo with the DOI *<u>10.5281/zenodo.15274014</u>* for 3-months visium spatial data and the DOI *<u>10.5281/zenodo.14988055</u>* for 20/23-months visium and snRNA data. The code, including helper objects and functions, for analysis and figure generation is available on GitHub: https://github.com/DanzerLab/PD_VisiumSpatial_MouseBrain_Aging. Human snRNA data from Smajic et al., 2022 can be accessed at GEO under <u>GSE157783</u>, data from Martirosyan et al., 2024 can be accessed at GEO under <u>GSE243639</u>, and data from Wang et al., 2022 can be accessed at GEO under <u>GSE19950</u>.

## Conflict of Interest

The authors declare no competing interests.

## Acknowledgements

We thank R. Bück for their excellent work and technical support.

## Funding Source

This work was supported and funded by the Deutsche Forschungsgemeinschaft (DFG) Emmy Noether Research Group DA 1657/2-1, SFB 1149, and SFB 1506 (Aging at Interfaces).

## Author Contribution

K.M. D. and J. LB. designed the project. V. B., A. G., J.K. K., L. M., and D. R. performed experiments and analyzed the data. V. B., A. G., and J. LB. prepared graphs and designed figures. E. G. performed quality control and annotation of cell types for snRNA-seq data. J. LB. performed all other bioinformatic analysis. V. G. and L. D. provided intellectual input. J. LB. and K.M. D. wrote the manuscript. All authors have revised or critically reviewed the article.

## Additional Data

Additional File 1.pdf: **Quality control of visium spatial of 20-months mice**

**a,** We checked for motoric deficits in our PD mouse model by measuring time on the rotarod. Using 10 control (OFF) and 9 PD (ON) mice, their time on the rotarod after training was measured between 13-14 months and 16-17 months of age. Only at the older time point did the ON mice show a significant motoric deficit compared to the OFF mice. The statistical testing was done by using a linear mixed-effects model of the condition x age effect with sex as a covariate and sample as a random intercept with significance set at a p value lower than 0.05. **b,** Scatter plot shows the ratio between unique number of detected genes (nGene) and unique molecular identifiers (nUMI). Each dot represents one visium spatial spot. Every spot that was kept after QC is marked red. **c**, The plot further supports the comparability between conditions as the overall distribution of nUMI peak for both conditions at around 12,000. **d,** After harmony integration and clustering the cluster 1, consisting of our region of interest the “Spreading Area” and the most similar region to our region of interest, was subclustered. We observed that integration enabled us to assign computationally the subcluster 2 as being representative of the “Spreading Area”. All other subclusters were used as “No Spreading Area”.

Additional File 2.pdf: **Annotation of 20-months visium spatial samples**

**a,** Left figure shows a schematic UMAP visualization of annotated clusters. The annotated regions reflect the anatomical characteristics based on their similarity, which can be seen on the right dendrogram. **b,** Furthermore, we were able to assign a canonical regional marker from mousebrain (mousebrain.org)(26) to each region. **c,** We observed some region to region differences in the nUMI but this mostly reflected their anatomical characteristics. Moreover, the differences were low enough that we deemed it as non-influential on our observations. **d**, Comparison of the final annotation of our visium spatial slides to the most similar slice from the Allen Mouse Brain Atlas(27, 28). Our annotations reflected known anatomical regions and the “Spreading Region” was part of the Substantia Nigra.

Additional File 3.xlsx: **Regional annotation markers found by wilcoxon rank sum test**

Markers found for each annotated region by using the FindMarkers function from Seurat(24). Only some exemplary markers were then used for visualization.

Additional File 4.xlsx: **Differential analysis results between OFF20M “Spreading Area” vs OFF20M “No Spreading Area”**

All 100 iterations of the differential analysis. Columns are named xxx-seed with the seed used for that specific iteration. Last 10 columns show the number of NA values per row, how many time a gene was significant (FDR < 0.05) and mean or median of log2 fold change, stat, p-value, and FDR.

Additional File 5.xlsx: **Differential analysis results between ON20M “Spreading Area” vs ON20M “No Spreading Area”**

All 100 iterations of the differential analysis. Columns are named xxx-seed with the seed used for that specific iteration. Last 10 columns show the number of NA values per row, how many time a gene was significant (FDR < 0.05) and mean or median of log2 fold change, stat, p-value, and FDR.

Additional File 6.pdf: **Region to region differences affect differential results and “Spreading Signature” changes is driven by the “Spreading Area”**

**a,** Heatmap of the expression levels of all significant (p.adj < 0.05) genes that were shared between ON20M and OFF20M across cell types. We see that a majority of those shared DEGs are cell type specific and are probably due to region to region differences in cell type composition. Thus, we only focused only on the ON20M specific DEGs by removing all shared DEGs between OFF20M and ON20M from the “Spreading Signature”. **b,** We further observe that the specific significance in ON20M and not in OFF20M for the final “Spreading Signature” comes from a change in the “Spreading Area” between conditions and not from “No Spreading Area”. **c,** We compared the log2 fold changes of “Spreading Signature” between all regions and “No Spreading Area” and observed that the difference was correlated to the cell type composition differences between regions. This means that α-syn oligomers affect the transcriptional landscape of regions differently due to their differences in cell type composition.

Additional File 7.xlsx: **“Spreading Signature” with functional and network information**

Every gene in the “Spreading Signature” with information about their function, the source for the function information, regulation direction in the “Spreading Area” to “No Spreading Area”, and network information (presence, which subnetwork, betweenness centrality, counted as central).

Additional File 8.xlsx: **Ligand receptor results from stlearn**

Ligand receptor results that was used for visualization from stlearn using their default ligand-receptor database

Additional File 9.pdf: **Spot deconvolution using CARD and changes in cell type composition**

**a,** Exemplary result plots of spot deconvolution by CARD based on our snRNA dataset. Cell type compositions reflected expected ratios based on anatomical characteristics. **b,** Changes of cell type composition in the “Spreading Area” from OFF20M to ON20M for cell types that were not in top 3 most changed cell types. Only cell type to show a significant pattern of downregulation was microglia. All other cell types obtained only miniscule differences.

Additional File 10.pdf: **Manual spatial annotation of Substantia Nigra using CLSM images**

**a,** Using sagittal brain slices of ON20M mice we stained for dopaminergic Neurons (TH) and V1S/SV2 (GFP). **b,** This enabled a clear manual annotation for Substantia nigra and its subparts Pars Compacta and Pars Reticulata.

Additional File 11.pdf: **Single nucleus RNA dataset quality control and annotation**

**a,** Annotation and similarity dendrogram based on harmony embeddings of our snRNA dataset. The dendrogram show a clear distinction between neuronal and non-neuronal cell types. **b,** Dot plot of a representative marker gene for each cell type. **c,** Cell type marker dot plot but for only Vglut1 neuron subtypes. **d,** With subclustering of astrocytes it was possible to get a distinction between *Slc1a2* enriched (glutamatergic) and *Slc6a11* enriched (GABAergic) astrocytes. **e,** Comparison of each cell type by their number of significant DEGs between OFF23M and ON23M and their summed nUMI count. Grey area represents the 95% confidence interval. Astro-Slc1a2 had many more DEGs compared to Astro-Slc6a11.

Additional File 12.xlsx: **Used markers for snRNA-seq annotation**

All used markers for snRNA-seq annotation including markers for subtypes of astrocytes, glutamatergic neurons and GABAergic neurons.

Additional File 13.xlsx: **Differential analysis results from Astro-Slc1a2 ON vs OFF**

All results from DESeq2 based differential analysis of 23-months snRNA-seq samples. ON vs OFF was tested using Wald testing. See methods for details.

Additional File 14.pdf: **Visium Spatial samples after QC and integration are comparable at 3M**

**a,** Left scatter plot shows the ratio between unique number of detected genes (nGene) and unique molecular identifiers (nUMI). Each dot represents one Visium Spatial spot. Every spot that was kept after QC is marked red. The right plot shows the nUMI across the annotated regions. **b,** After harmony integration and clustering the cluster 1, consisting of our region of interest the “Spreading Area” and the most similar region to our region of interest, was subclustered. We observed that integration enabled us to assign computationally the subcluster 2 as being representative of the “Spreading Area”. All other subclusters were used as “No Spreading Area”. **c,** “Spreading Area” and “No Spreading Area” showed the same difference between mRNA and protein expression of V1S/SV2 in the 3M mice as in the 20M mice.

Additional File 15.pdf: **snRNA results also point to astrocytic dysfunction in the glutamate cycle**

**a,** Venn diagram showing intersection between the “Spreading Signatures” at 3M and 20M. **b,** Network built from the DEGs between snRNA OFF23M and ON23M in Astro-Slc1a2 and expression data of ON23M. The top 10 central genes and the two shared genes also points to astrocytic dysfunction in the glutamate cycle at the synapses.

Additional File 16.xlsx: **Differential analysis results between OFF3M “Spreading Area” vs OFF3M “No Spreading Area”**

All 100 iterations of the differential analysis. Columns are named xxx-seed with the seed used for that specific iteration. Last 10 columns show the number of NA values per row, how many time a gene was significant (FDR < 0.05) and mean or median of log2 fold change, stat, p-value, and FDR.

Additional File 17.xlsx: **Differential analysis results between ON3M “Spreading Area” vs ON3M “No Spreading Area”**

All 100 iterations of the differential analysis. Columns are named xxx-seed with the seed used for that specific iteration. Last 10 columns show the number of NA values per row, how many time a gene was significant (FDR < 0.05) and mean or median of log2 fold change, stat, p-value, and FDR.

Additional File 18.xlsx: **Significant differential genes from Astro-Slc1a2 analysis with functional and network information**

Every gene with genes deemed central in the network marked in red. Every central gene has information about their function, the source for the function information, regulation direction, and network information (presence, betweenness centrality, counted as central).

Additional File 19.xlsx: **Differential analysis results from Vglut1-Col19a1 ON vs OFF**

All results from DESeq2 based differential analysis of 23-months snRNA-seq samples. ON vs OFF was tested using Wald testing. See methods for details.

Additional File 20.xlsx: **Differential analysis results from Vglut1-Npas3 ON vs OFF**

All results from DESeq2 based differential analysis of 23-months snRNA-seq samples. ON vs OFF was tested using Wald testing. See methods for details.

Additional File 21.xlsx: **Relevant significantly differential genes from ON vs OFF comparisons in Vglut1-Npas3 and Vglut1-Col19a1**

Every relevant gene has information about their function, the source for the function information, and regulation direction.

Additional File 22.xlsx: **Final list of genes deemed central to Astro-GluN dysfunctions**

List of genes deemed central to our proposed model of Astro-GluN dysfunctions in the progression of α-syn oligomer pathology. Each row has information about one gene’s function, observed regulation direction, source of the function information, and on which data the gene was deemed central.

## Notes

https://github.com/DanzerLab/PD_VisiumSpatial_MouseBrain_Aging

https://zenodo.org/records/15274014

https://zenodo.org/records/14988055

### Competing Interest Statement

The authors have declared no competing interest.

### Summary of Updates

Multiple Figures were updated for clarity and understanding. Manuscript text were also edited for clarity and a Limitation of the Study was added.

