## Supplementary figures and images for "Spreading alpha-synuclein Oligomers Trigger Astrocyte Changes and Astrocyte-Glutamatergic Neuron system dysfunction in an Age-related Manner"

### Suppl. Figure 1

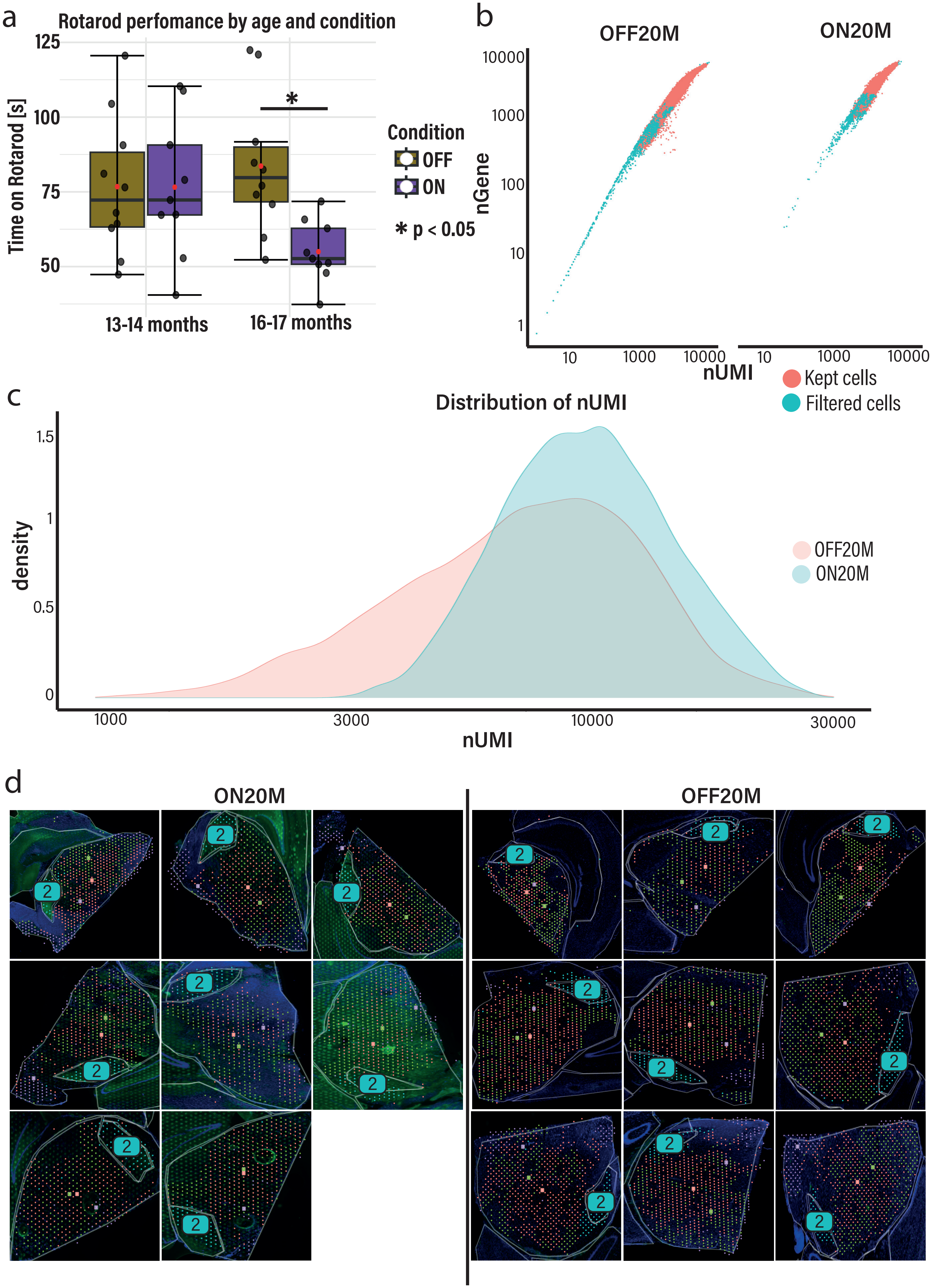

### Suppl. Figure 2

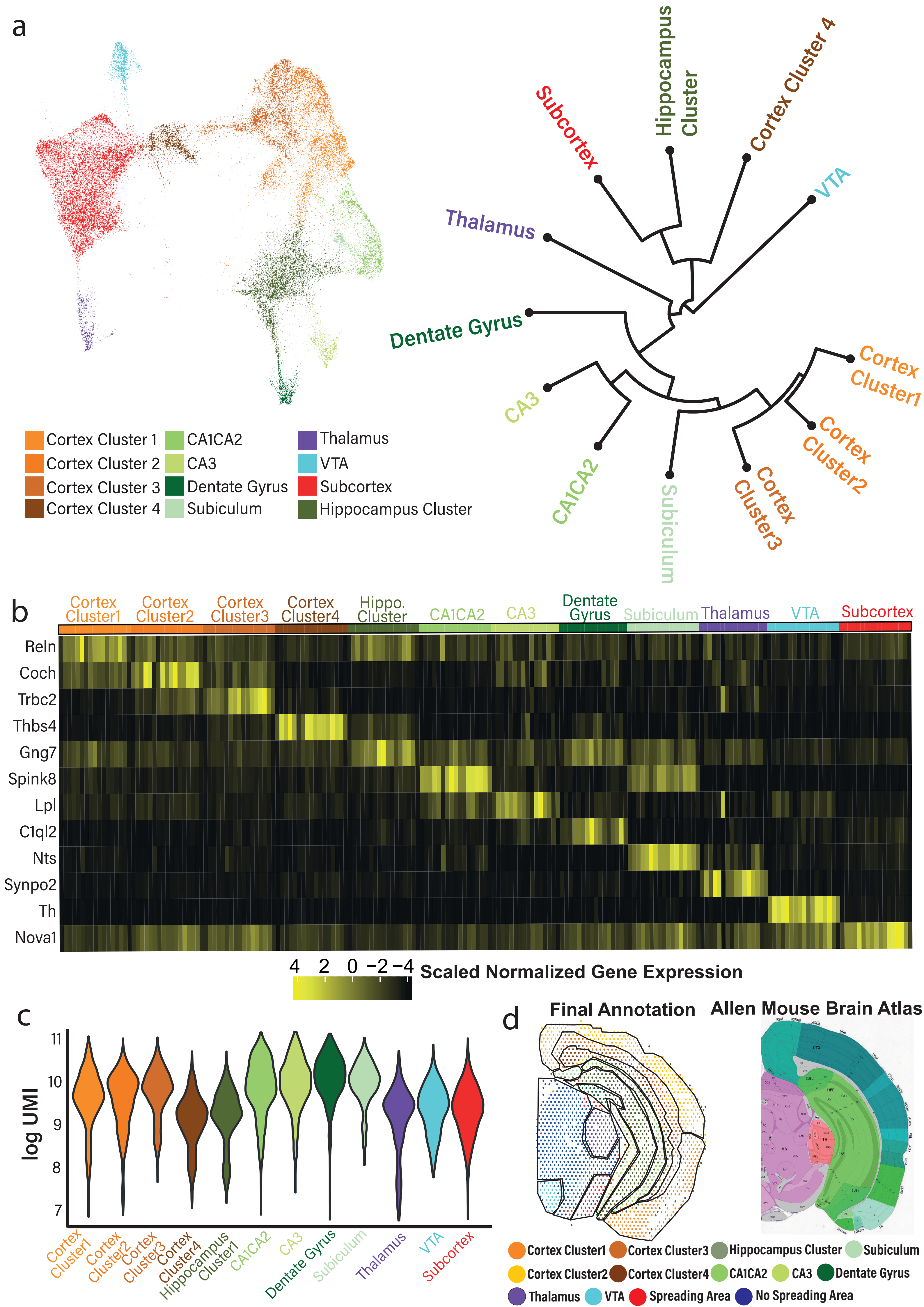

### Suppl. Figure 3

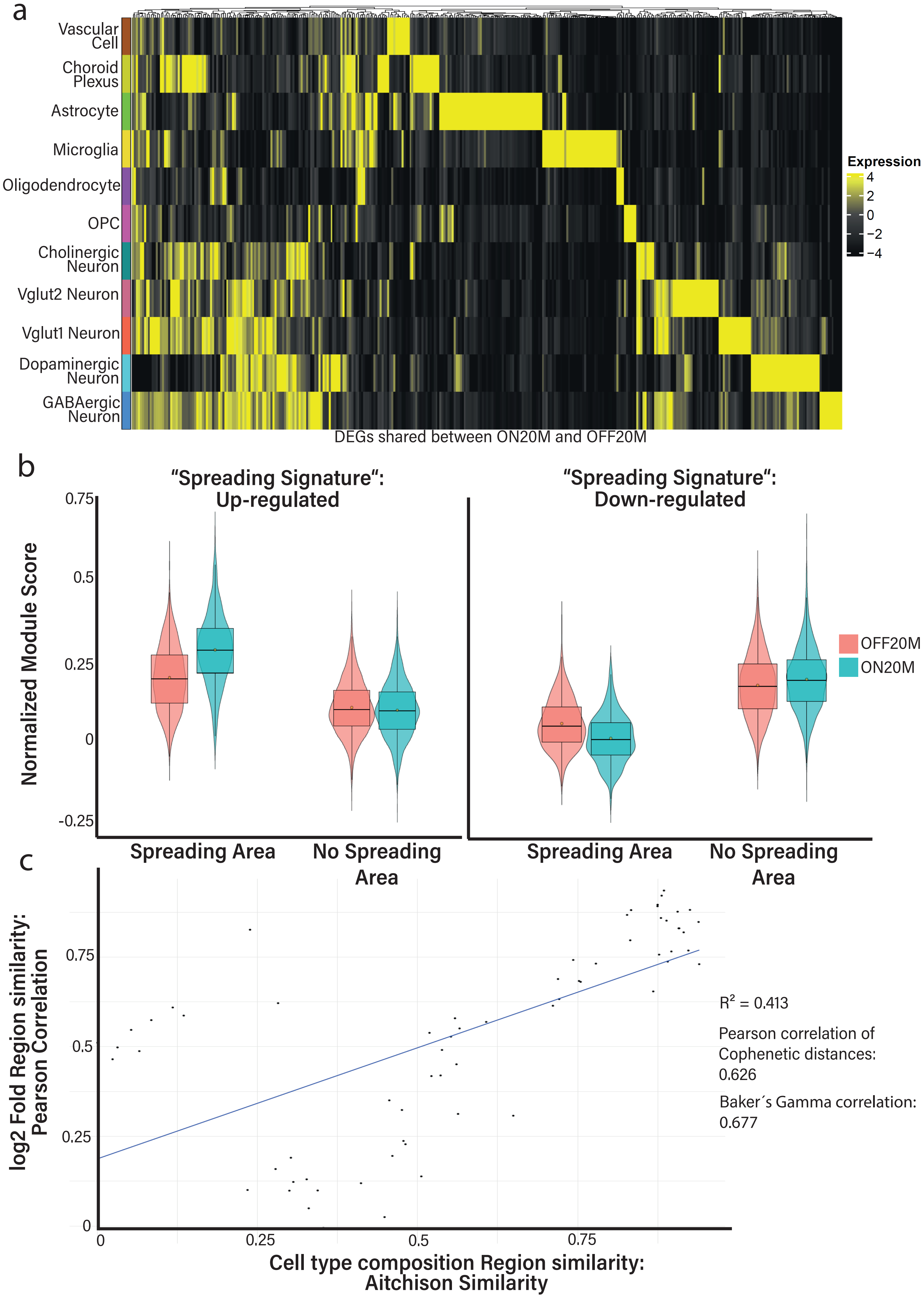

### Suppl. Figure 4

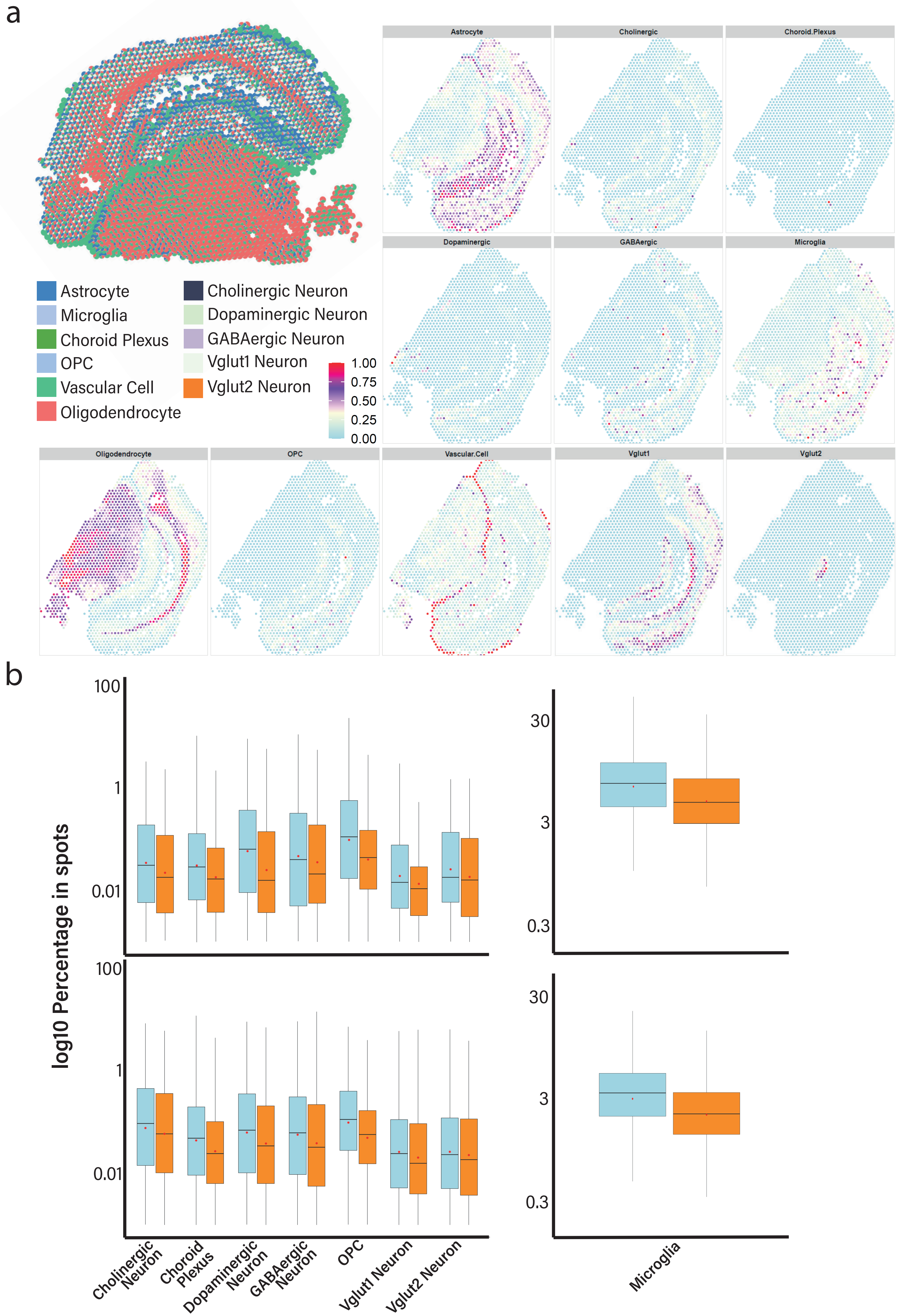

### Suppl. Figure 5

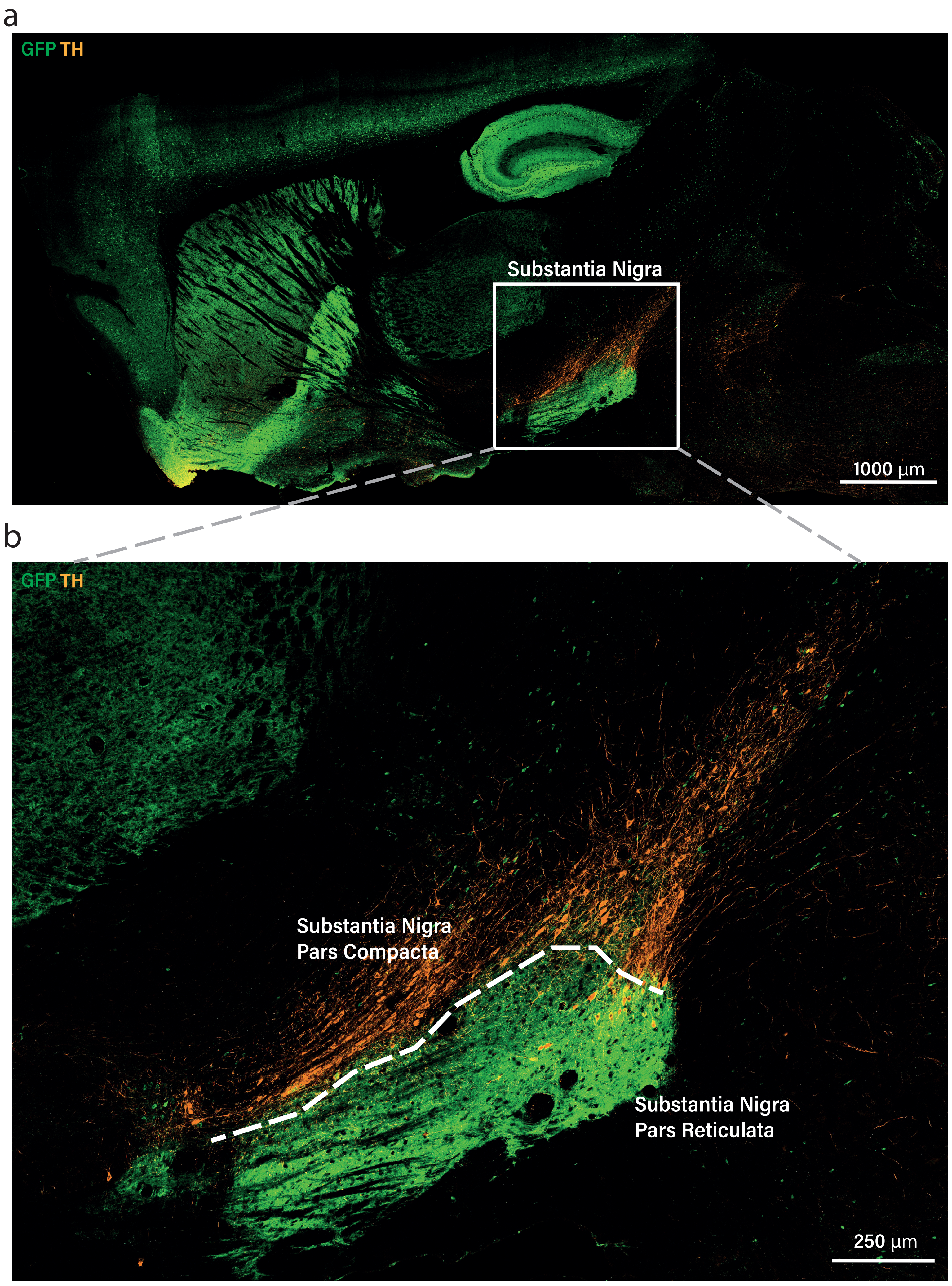

### Suppl. Figure 6

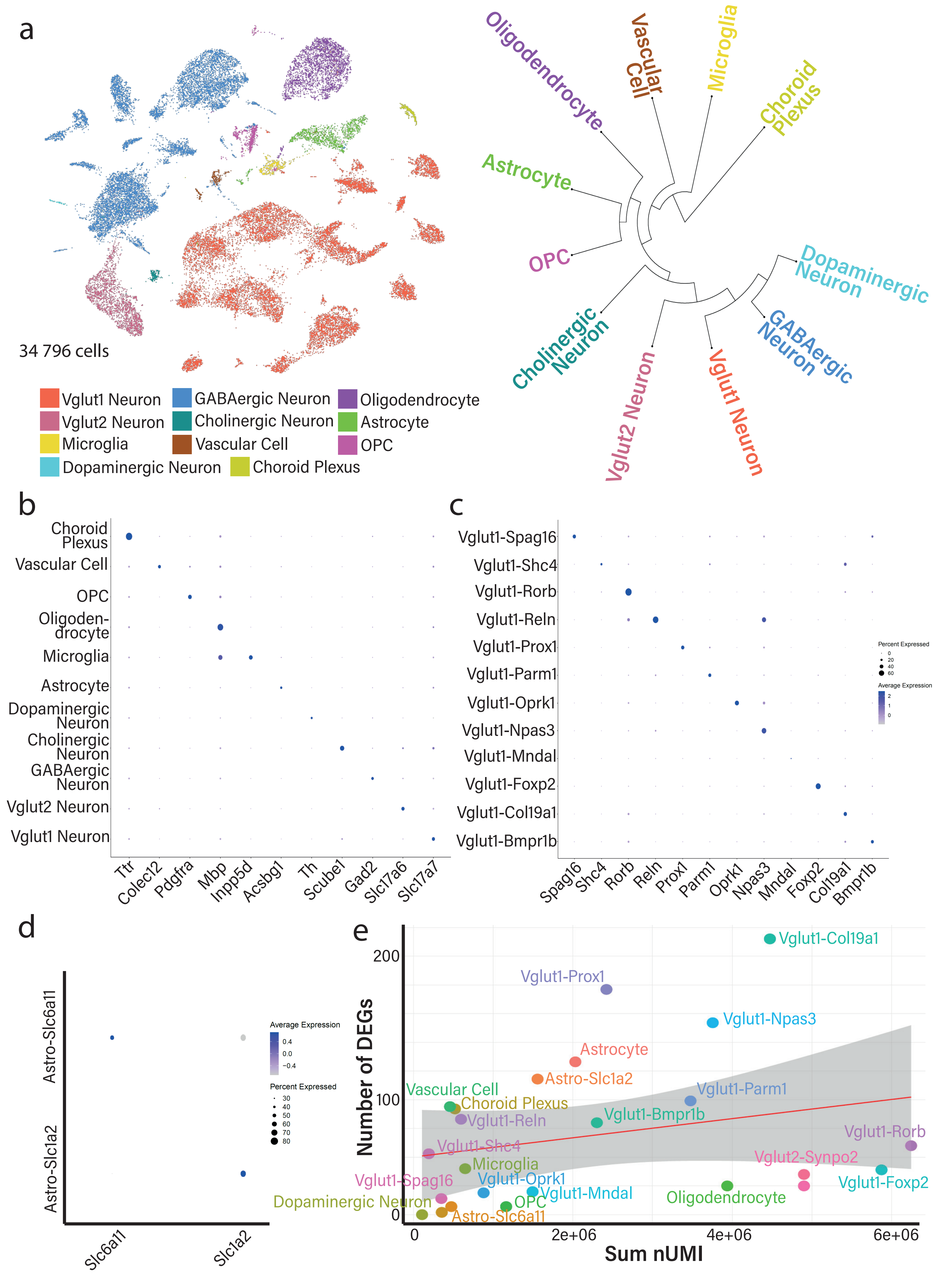

### Suppl. Figure 7

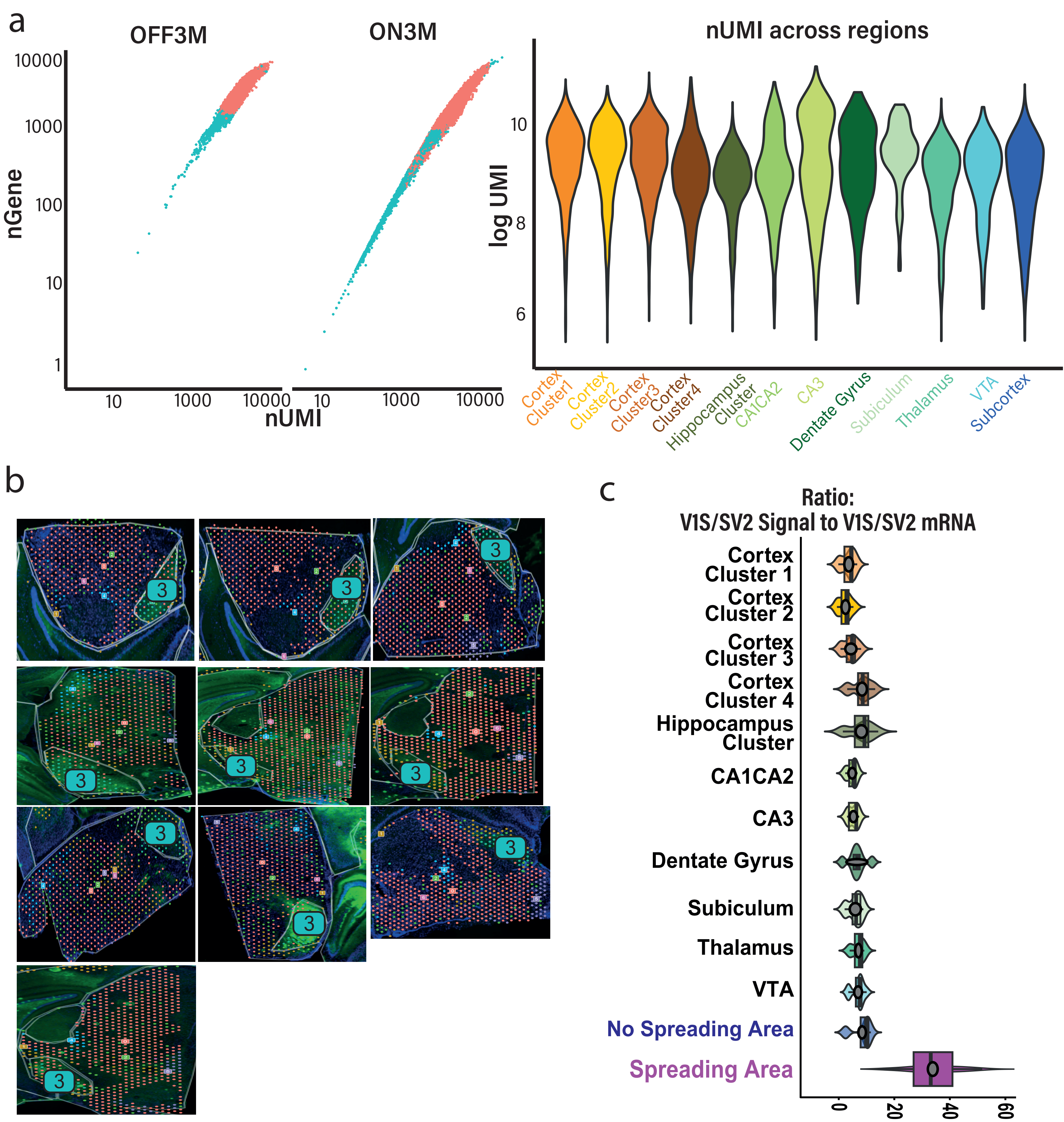

### Suppl. Figure 8

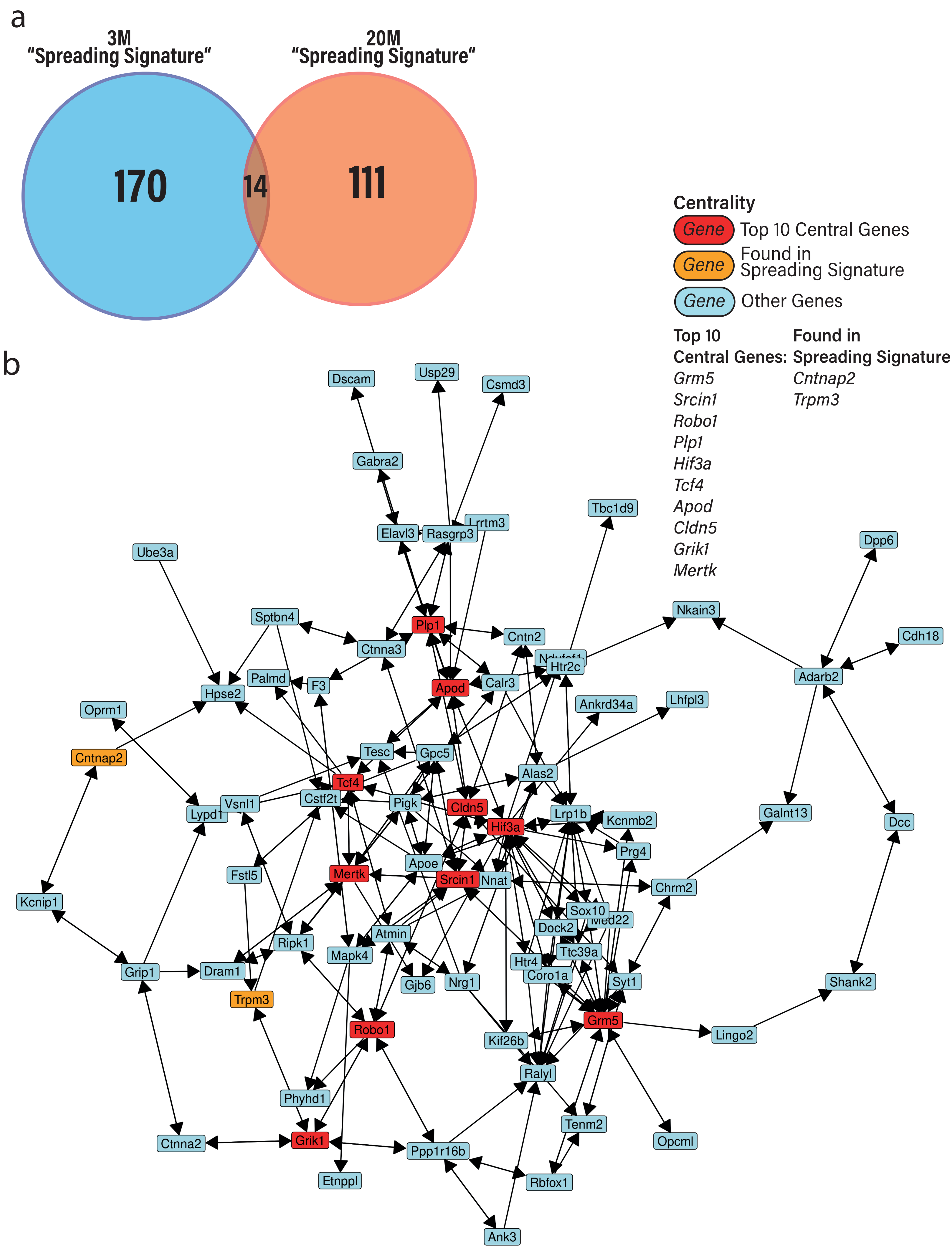
